# Regulation of membrane homeostasis by TMC1 mechanoelectrical transduction channels is essential for hearing

**DOI:** 10.1101/2021.09.24.461722

**Authors:** Angela Ballesteros, Kenton J. Swartz

## Abstract

The mechanoelectrical transduction (MET) channel complex of auditory hair cells converts sound into electrical signals, allowing us to hear. After decades of research, the transmembrane-like channel 1 and 2 (TMC1 and TMC2) have been recently identified as pore-forming subunits of the MET channels, but the molecular peculiarity that differentiates these two proteins and makes TMC1 essential for hearing remains elusive. Here, we show that TMC1, but not TMC2, is essential for membrane remodeling triggered by a decrease in intracellular calcium concentration. We demonstrate that inhibition of MET channels or buffering of intracellular calcium lead to pronounced phosphatidylserine externalization, membrane blebbing and ectosome release at the hair cell sensory organelle, culminating in the loss of TMC1 protein. Moreover, three TMC1 deafness-causing mutations cause constitutive phosphatidylserine externalization that correlates with the deafness phenotype, suggesting that the mechanisms of hearing loss involve alterations in membrane homeostasis.

## INTRODUCTION

Auditory and vestibular hair cells of the inner ear are the sensory cells responsible for perceiving sound and head movements, respectively. The mechanosensory apparatus consists of a bundle of 50-100 specialized actin-based cilia, termed stereocilia, at the apical region of hair cells (Fettiplace and Kim, 2014). In mammals, stereocilia are arranged in 3 rows of increasing height interconnected by several protein filaments (Furness and Hackney, 2006), including the tip link that transfers mechanical stimuli to the mechanoelectrical transduction (MET) channels (Assad et al., 1991). After sound stimulation, the stereocilia are deflected toward the tallest stereocilia row, opening the MET channel and leading to depolarization of the hair cell and conversion of the mechanical stimulus into an electrical response (Hudspeth and Corey, 1977).

Decades of research have led to the identification of the transmembrane-like channel 1 and 2 proteins (TMC1 and TMC2) as pore-forming subunits of the MET channel (Zheng and Holt, 2021, Kawashima et al., 2015). Mammalian auditory hair cells express both TMC1 and TMC2, but their expression patterns vary during development, and they confer different properties to the MET channel (Corns et al., 2017, Pan et al., 2013, Kawashima et al., 2011). Importantly, expression of TMC2 is insufficient for maintaining normal hearing in mice lacking TMC1 (Nakanishi et al., 2018), suggesting that TMC1 must have additional functions required for hearing. More than 30 autosomal dominant (DFNA36) and recessive (DFNB7/11) mutations in the *tmc1* gene cause deafness in humans, highlighting the importance of TMC1 in sound transduction (Kawashima et al., 2015, Kurima et al., 2002). In contrast, no deafness-causing mutations have been identified in *tmc2*. The dominant M412K and D569N, and the recessive D528N TMC1 mutations have been extensively characterized in mice (Beurg et al., 2015, Vreugde et al., 2002, Beurg et al., 2019, Beurg et al., 2021, Marcotti et al., 2006, Pan et al., 2013, Corns et al., 2016). The effects of these three mutations are more severe in homozygous than in heterozygous mice and they all decrease the calcium (Ca^2+^) permeability of the MET channel. However, since recessive D528N generally alters the MET channel properties more severely than dominant M412K and D569N, the deafness phenotype of these mutations does not appear to relate to the functional properties of the MET channel. Thus, the molecular mechanisms of TMC1-related deafness remain enigmatic.

We and others have previously reported that TMC proteins are evolutionarily and structurally related to TMEM16, a family of Ca^2+^-activated chloride channels, some of which can also function as lipid scramblases (Ballesteros et al., 2018, Pan et al., 2018, Medrano-Soto et al., 2018, Kunzelmann et al., 2016). Lipid scrambling alters the phospholipid bilayer asymmetry of the mammalian plasma membrane resulting in the externalization of phosphatidylserine (PS), a hallmark of many relevant biological processes such as apoptosis, blood coagulation, myoblast fusion, fertilization, synapse pruning, or photoreceptor disc shedding (Bevers and Williamson, 2016). Inspired by this TMC-TMEM16 structural relation, we investigated a potential role of TMC proteins in the regulation of hair cell membrane homeostasis using Airyscan super-resolution confocal microscopy and 13 different transgenic mouse lines. We found that pharmacological inhibition of the MET channel, breakage of the tip links, and buffering of intracellular Ca^2+^ trigger PS externalization and the formation of PS-positive membrane blebs at the stereocilia and apical hair cell region in a TMC1-dependent and TMC2-independent manner. Furthermore, we report that hair cells impaired by TMC1 deafness-causing mutations display constitutive PS externalization, emphasizing the importance of membrane homeostasis for hearing. Importantly, our findings show that TMC1 dominant and recessive mutations alter PS externalization by different mechanisms involving intracellular Ca^2+^, revealing that while M412K and D569N are gain-of-function mutations, D528N is a loss-of-function mutation. In short, we unveil a novel role for TMC1 in the regulation of hair cell membrane homeostasis that is essential for hearing.

## RESULTS

### Blockage of MET channels triggers PS externalization and membrane blebbing at the apical hair cell region

Rapid PS externalization and membrane blebs have been previously observed at the apical region of hair cells treated with ototoxic aminoglycoside antibiotics, which are well-known inhibitors of the MET channel (Goodyear et al., 2008, Richardson and Russell, 1991). To examine whether this dysregulation of membrane homeostasis is caused by the blockage of the MET current at rest, we examined the induction of PS externalization and membrane blebbing in hair cells treated with the aminoglycoside neomycin, and with 2 other non-ototoxic and well-established MET channel blockers, benzamil and curare (Glowatzki et al., 1997, Rusch et al., 1994, Kroese et al., 1989). For this purpose, externalized PS at the cell membrane was detected using the PS specific binding protein annexin V (AnV) and the overall hair cell surface was visualized by staining with fluorescently labeled wheat germ agglutinin (WGA), which does not affect MET currents (Meers and Mealy, 1993, Richardson et al., 1989). To visualize the membrane, we performed similar experiments in mT/mG^Tg/+^ mice expressing one copy of the membrane-targeted tandem dimer tomato (mtdTomato) fluorescent reporter, which present unaltered MET (Ballesteros et al., 2021).

As shown in Figure 1, untreated cochlear hair cells from 6 days old (P6) mice show no specific labeling of AnV, indicating that PS is not exposed on these cells. However, 25 min incubation with 100 μM neomycin, benzamil, or the less permeable antagonist curare (Farris et al., 2004, Glowatzki et al., 1997, Rusch et al., 1994) triggered robust PS externalization at the apical hair cells region in wild type (Figure 1A-B) and mT/mG^Tg/+^ mice (Figure 1C-D). Neomycin and benzamil induced PS externalization in both outer (OHCs) and inner hair cells (IHCs), whereas curare-induced PS externalization was mainly restricted to OHCs. Although, the MET IC50 of curare was estimated in murine OHCs and higher curare concentrations may be needed to fully block the MET channels in IHCs (Glowatzki et al., 1997), these data demonstrate that chemically distinct MET inhibitors trigger PS externalization.

**FIGURE 1.**
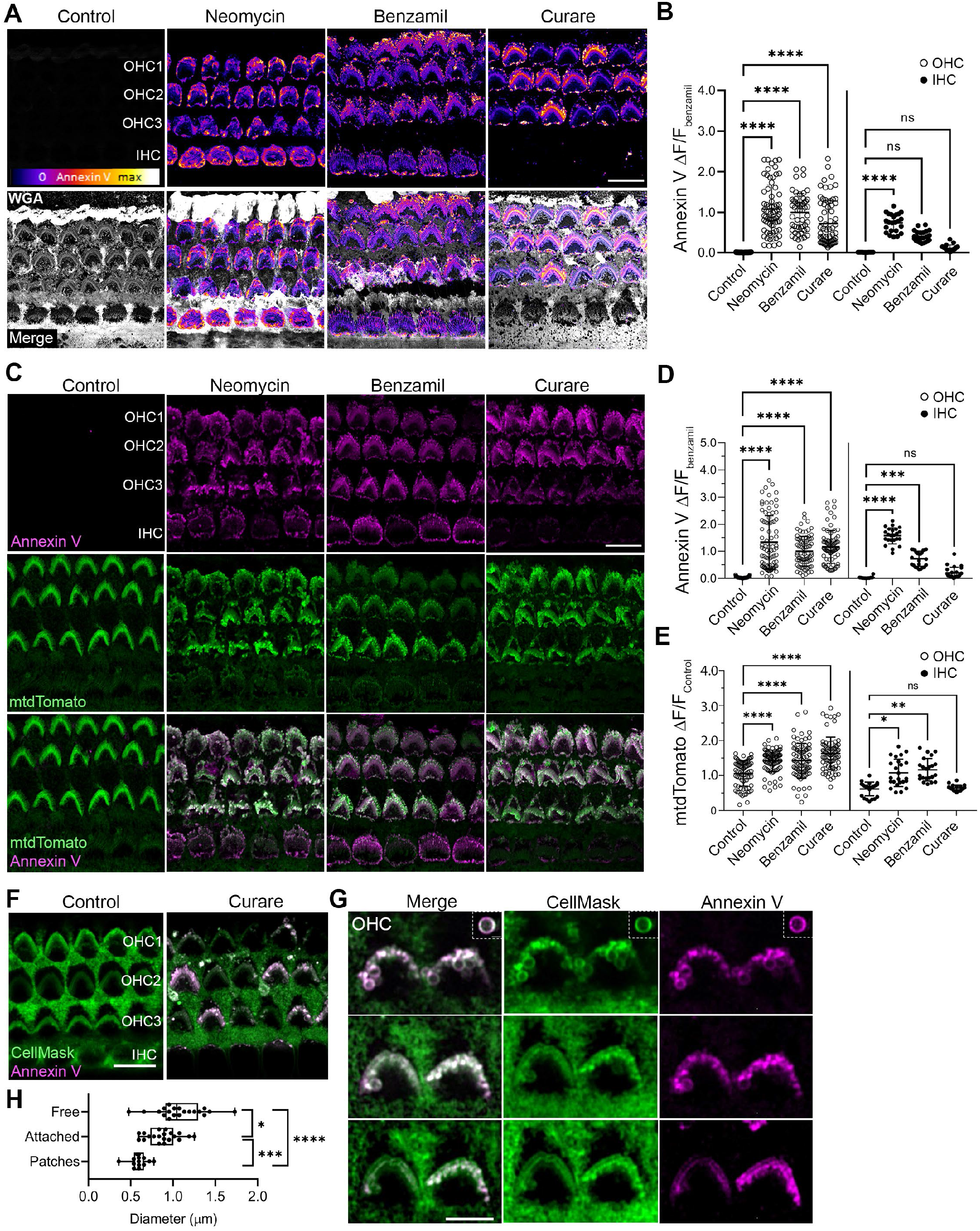
Blockade of the MET channel triggers PS externalization and membrane blebbing. **A**) Confocal images of P6 wild type hair cells incubated with WGA (gray) and AnV in the absence (control) or presence of 100 μM neomycin, benzamil or curare. **B)** Quantification of AnV fluorescence intensity in IHCs (○) or OHCs (●) treated as in A. **C)** Confocal images of P6 mT/mG^Tg/+^ hair cells expressing the mtdTomato membrane reporter (green) treated as in A. Quantification of the AnV (**D**) or mtdTomato (**E**) fluorescence intensity in IHCs and OHCs treated as in C. **F)** Live imaging of wild type hair cells labeled with CellMask (green) and AnV (magenta) in the absence (control) or presence of curare. **G)** Confocal planes of OHCs treated with curare as in F. **H**) Diameter of AnV positive vesicles (n=20). In B, D and E, mean ± SD is shown for n = 85-70 OHCs and 27-22 IHCs from 2 cochleae. One-way ANOVA analysis was performed (n.s. p>0.05, \**p*<0.05, \*\**p*<0.01, \*\*\**p*<0.001, and \*\*\*\**p*<0.0001). Scale bar is 10 μm in A, C and F, and 5 μm in G. See also Figure S1 and S2.

In addition to PS externalization, neomycin is known to induce membrane blebbing and increase hair cell capacitance, indicating that PS externalization is accompanied by membrane addition to the hair cell surface (Goodyear et al., 2008, Forge and Richardson, 1993). Consistent with this, we found that while the mtdTomato reporter labeled the membrane of untreated hair cells isolated from mT/mG^Tg/+^ mice homogenously, it was enriched in vesicle-like structures and membrane blebs in hair cells treated with neomycin, benzamil or curare (Figure 1C and S1). In addition, we observed an increase in the mtdTomato fluorescence intensity in OHCs and IHCs treated with neomycin or benzamil and in OHCs treated with curare (Figure 1E), which parallels the changes in AnV signal (Figure 1D). These data indicate that inhibition of the MET channel leads to membrane addition and blebbing at the apical region of hair cells.

The WGA labeling and the resolution provided by the Airyscan LSM880 confocal microscope allowed us to pinpoint the externalized PS. Neomycin is known to induce PS externalization at the stereocilia, around the apical surface of the hair cell, and at the kinocilial area (Goodyear et al., 2008). Accordingly, we observed that treatment with benzamil, neomycin, or curare led to PS externalization at these same regions. Hair cells treated with neomycin presented larger AnV-positive patches around the hair cells and at the kinocilial area than the other treatments (Figure S1 and S2, blue arrows). These AnV-positive patches or vesicles were also enriched with mtdTomato, suggesting that PS externalization and membrane blebbing are related. Interestingly, we observed that externalized PS and mtdTomato accumulate at the tips of the shorter stereocilia rows (Figure S1 and S2, pink arrows), where TMC1, TMC2 and the MET channel are located (Beurg et al., 2009, Kurima et al., 2015). Vesicle-like particles labeled with WGA, AnV and mtdTomato were observed bulging at the tips of the stereocilia and around the hair cell surface (Figure S1 and S2, yellow arrows), indicating outward budding and vesiculation of the membrane.

We also confirmed the formation of these AnV-positive patches, blebs and vesicles upon blockage of the MET in live-cultured wild type hair cells labeled with the membrane marker CellMask. AnV specifically detected externalized PS at the hair cell bundle of curare-treated cells but failed to detect PS in the untreated hair cells (Figure 1F). AnV and CellMask-positive patches, blebs and vesicles were observed at the tips of the stereocilia and floating near the stereocilia bundle in treated samples (Figure 1G). The average diameter of free vesicles (1.05 ± 0.27 μm) was significantly larger than that of stereocilia-attached vesicles (0.55 ± 0.28 μm), which suggests that these vesicles are ectosomes shedding from the stereocilia membrane (Cocucci and Meldolesi, 2015). We conclude that pharmacological inhibition of the MET channel leads to PS externalization, membrane blebbing and ectosome release in the apical region of murine auditory hair cells, suggesting that blockage of the MET channel is accompanied by dysregulation of the apical hair cell membrane homeostasis.

### Tip link disruption leads to PS externalization and membrane blebbing at the apical hair cell region

We next explored whether manipulations that mechanically disturb the MET channel current would also trigger PS externalization. For this, we analyzed PS externalization and membrane blebbing in hair cells from P6 mT/mG^Tg/+^ mice treated with Hank’s balanced salt solution (HBSS, control) or 5 mM BAPTA in Ca^2+^ and Mg^2+^ free HBSS (HBSS-CFM), a condition known to break the tip links that mechanically gate the MET channel (Indzhykulian et al., 2013, Assad et al., 1991). We found a robust PS externalization and accumulation of AnV and mtdTomato fluorescence signals at the stereocilia, vestigial kinociliary site and blebs in BAPTA-treated hair cells but not in the control (Figure 2A). In addition, AnV and mtdTomato fluorescence intensities were increased in BAPTA-treated hair cells (Figure 2B-C). Comparable results were obtained in hair cells from P6 wild type mice (Figure 2D-E). Importantly, we did not detect AnV labeling in hair cells incubated in HBSS-CFM alone (Figure 2D), indicating that the PS externalization observed in BAPTA-treated cells was not caused by the low cation concentration of this buffer. Interestingly, like our observations with the MET blockers, AnV staining was more robust in OHCs than in IHCs and only a few IHCs exhibited detectable externalized PS (Figure 2B and E). It is important to note that due to the Ca^2+^-dependent PS binding of AnV, cochlear explants were incubated for 5 min in HBSS containing AnV after BAPTA-treatment. Although the regeneration of the tip links takes hours (Zhao et al., 1996), we wanted to confirm these results using a recombinant C2 domain of lactadherin/MFG-E8 fused to the fluorescent protein clover (clover-Lact-C2), which binds PS in the absence of Ca^2+^ (Del Vecchio and Stahelin, 2018). Consistent with the AnV results, clover-Lact-C2 labeling was detected in BAPTA-treated cells but not in the control (Figure S3), further supporting the specific detection of externalized PS in response to tip link disruption. These data indicate that PS is externalized when the tip links no longer transduce mechanical force to open the MET channels.

**FIGURE 2.**
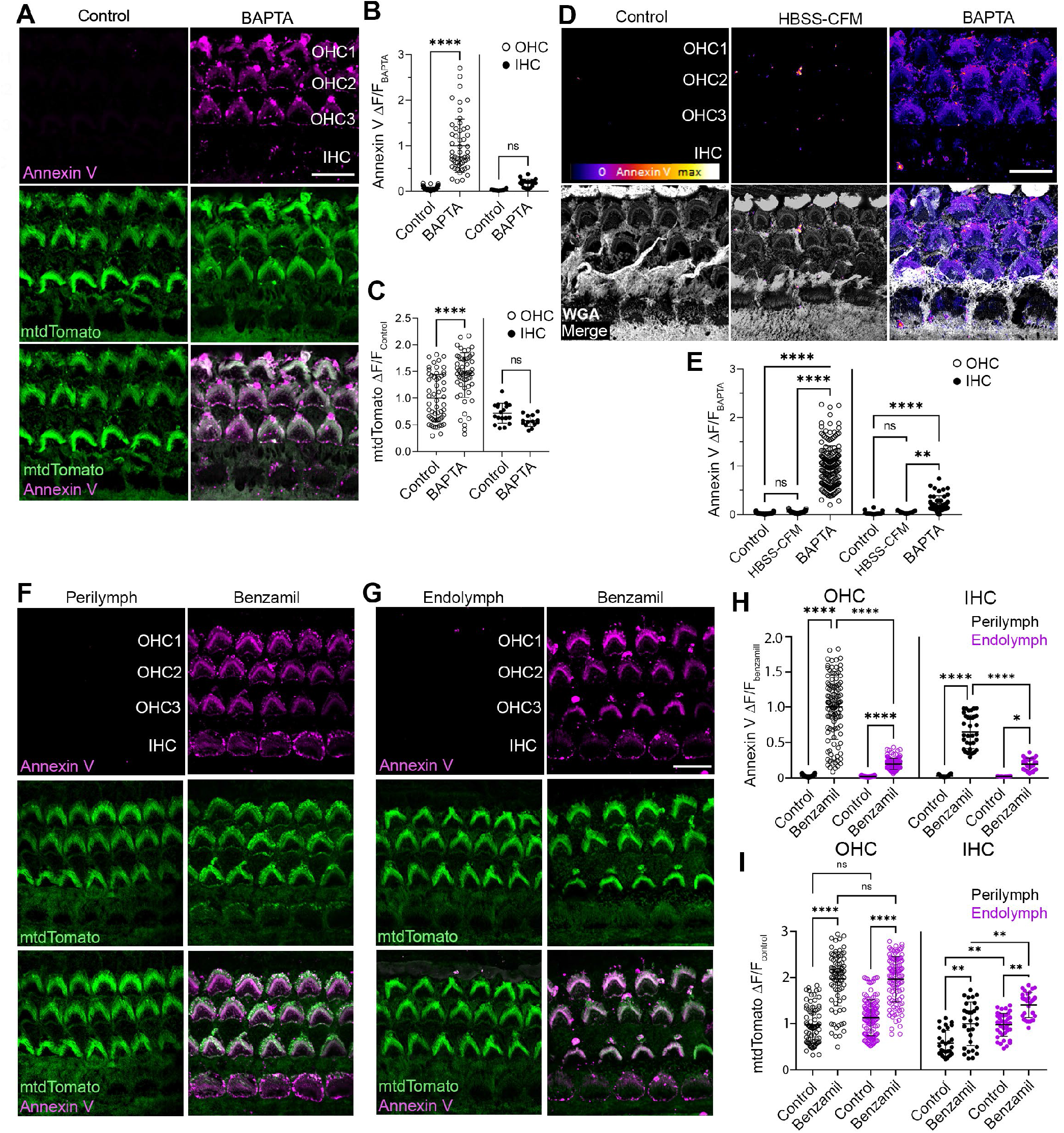
PS externalization and membrane blebbing after tip links disruption and in biological extracellular solutions. **A)** Confocal images of P6 mT/mG^Tg/+^ hair cells expressing mtdTomato (green) treated with HBSS (control) or 5 mM BAPTA before the addition of AnV (magenta). Quantification of the AnV (**B**) and mtdTomato (**C**) fluorescence intensity at the apical region of IHCs (○) and OHCs (●) treated as in A. **D**) Confocal images of wild type hair cells treated with HBSS (control), HBSS cation-free media (HBSS-CFM) or 5mM BAPTA in HBSS-CFM containing WGA (grey) before the addition of AnV (Fire LUT). **E**) AnV fluorescence intensity quantification as in D. **F)** Confocal images of P6 mT/mG^Tg/+^ mice hair cells untreated or treated with benzamil in perilymph-like buffer containing AnV. **G)** Confocal images of P6 mT/mG^Tg/+^ hair cells untreated or treated with benzamil in endolymph-like buffer containing AnV. Quantification of AnV (**H**) and mtdTomato (**I**) fluorescence intensity as in H and J. In B-C, E, H and I, mean fluorescence intensity ± SD is represented for n = 126-82 OHCs and 43-19 IHCs from 2 cochleae. One-way ANOVA analysis was performed (n.s. p>0.05, \*\**p*<0.01, and \*\*\*\**p*<0.0001). Scale bar is 10 μm. See also Figure S3.

### PS externalization and membrane remodeling after MET blockage occur in perilymph-like conditions

Most of our experiments were performed in HBSS containing 1.26 mM Ca^2+^. However, in the intact cochlea, the apical hair cell region is immersed in an endolymph solution rich in Na^+^ and K^+^ and low in Ca^2+^, whereas the basal region is bathed in perilymph with an ion composition similar to HBSS (Bosher and Warren, 1978). Thus, we next tested whether PS externalization occurs in a physiological environment by incubating hair cells from P6 mT/mG^Tg/+^ mice with AnV in perilymph (1.3 mM Ca^2+^) or endolymph buffer (25 μM Ca^2+^) in the absence or presence of 100 μM benzamil. In the absence of benzamil, we did not detect externalized PS in explants incubated in either perilymph or endolymph buffer, indicating that reduced extracellular Ca^2+^ concentrations alone do not trigger PS externalization (Figure 2F-H). This is consistent with our data with HBSS-CFM (Figure 2D-E). However, MET blockade with benzamil triggered PS externalization in hair cells immersed in either perilymph or endolymph buffer (Figure 2F-H). Interestingly, while the increase in mtdTomato fluorescence intensity was similar in benzamil treated samples, AnV signal in the endolymph buffer was lower than in the perilymph buffer (Figure 2H-I). Therefore, although extracellular Ca^2+^ influences the extent of PS externalization, these results indicate that PS externalization and membrane blebbing occur under physiological ionic conditions.

### Transgenic mice lacking functional MET channels fail to externalize PS and remodel the membrane

We have demonstrated that physical and pharmacological disruption of MET current trigger PS externalization and membrane blebbing, suggesting that functional MET channels are required for maintenance of membrane homeostasis. To test this hypothesis, we examined PS externalization in response to MET channel blockers in hair cells from TMC1 and TMC2 double knock-out mice (TMC1^-/-^ TMC2^-/-^), which lack functional MET channels (Kawashima et al., 2011). Furthermore, we generated a triple transgenic mT/mG^Tg/+^ TMC1^-/-^ TMC2^-/-^ mouse expressing the mtdTomato reporter to visualize the hair cell membrane. As previously reported (Beurg et al., 2018), we found that the morphology of TMC1^-/-^ TMC2^-/-^ hair cell bundles was altered; IHCs presenting additional stereocilia rows, and OHCs were arranged in a “U” shape (Figure S4E). Hair cells from mT/mG^Tg/+^ TMC1^-/-^ TMC2^-/-^ (Figure 3A-C) or TMC1^-/-^ TMC2^-/-^ mice (Figure S4A-D) were incubated with different MET blockers like in Figure 1. In contrast to our results with wild type hair cells, treatment with neomycin, benzamil, or curare failed to trigger PS externalization in hair cells from these 2 MET-lacking mouse lines (Figure 3A-C and Figure S4A and S4D). Furthermore, we did not observe any alteration of the apical hair cell membrane or an increase of the mtdTomato fluorescence intensity upon MET blockage in mT/mG^Tg/+^ TMC1^-/-^ TMC2^-/-^ hair cells (Figure 3A-C), suggesting that TMC1 and TMC2 are required for membrane remodeling after MET channel blockade. Additionally, we analyzed PS externalization after benzamil treatment in hair cells from P6 homozygous mT/mG mice (mT/mG^Tg/Tg^), which unlike their heterozygous counterparts, lack functional MET channels as indicated by their failure to uptake the MET channel-permeable FM1-43 dye (Ballesteros et al., 2021). As shown in Figure 3D-E, wild type and mT/mG^Tg/+^ hair cells externalized PS upon benzamil treatment whereas benzamil-treated hair cells from mT/mG^Tg/Tg^ littermates failed to externalize PS. Furthermore, mtdTomato fluorescence intensity increased after benzamil treatment in mT/mG^Tg/+^ hair cells but not in cells from mT/mG^Tg/Tg^ mice (Figure 3F). Thus, we conclude that TMC1 and TMC2 proteins or functional MET channels are required for PS externalization and apical hair cell membrane remodeling after MET inhibition.

**FIGURE 3.**
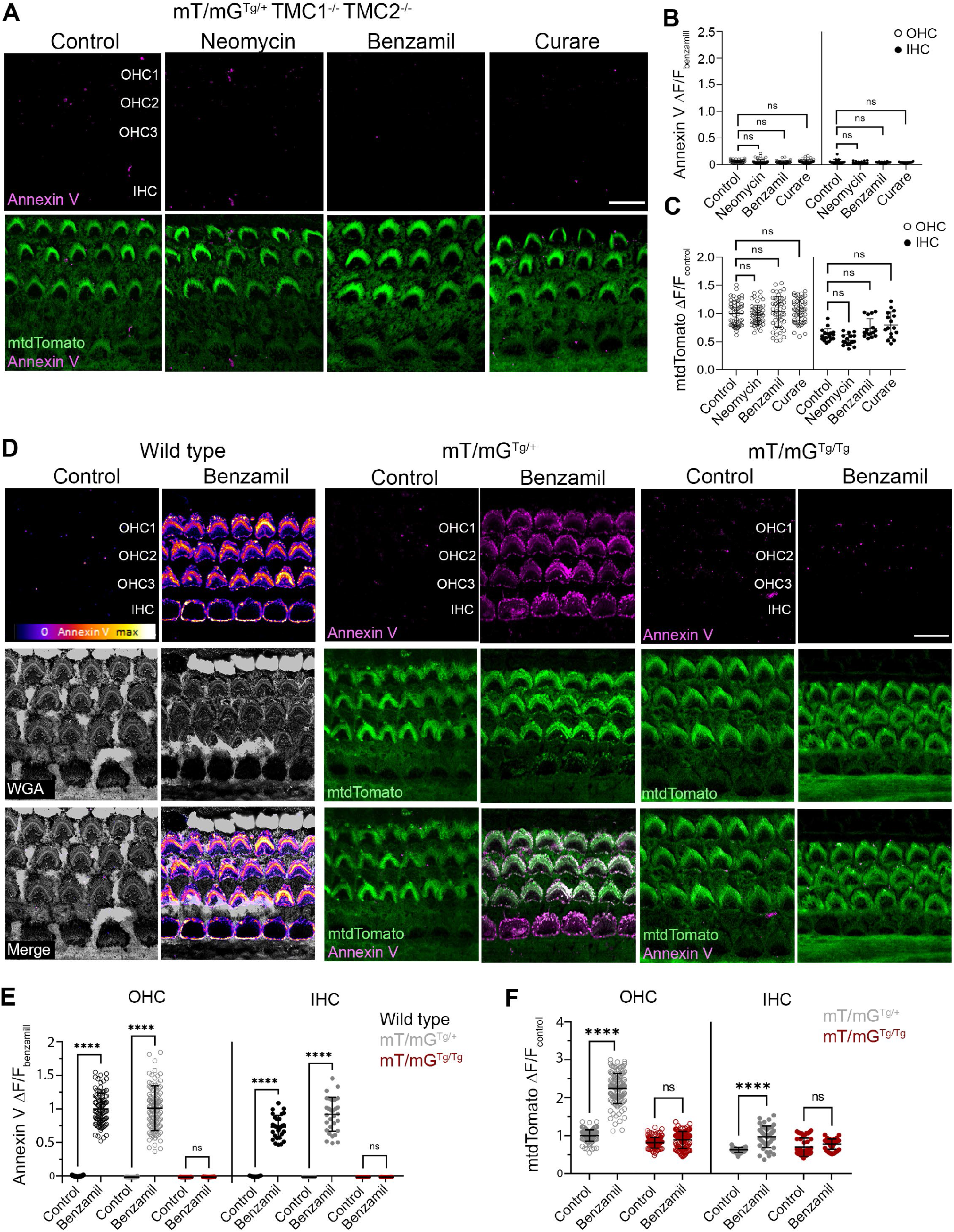
PS externalization and membrane blebbing in mice lacking MET. **A)** Confocal images of P6 mT/mG^Tg/+^ TMC1^-/-^ TMC2^-/-^ hair cells incubated with HBSS containing AnV in the absence (control) or the presence of 100 μM neomycin, benzamil or curare. Quantification of the AnV (**B**) and mtdTomato (**C**) fluorescence intensity at the apical region of the OHCs (○) and IHCs (●) treated as in A. **D)** Confocal images of hair cells from P6 wild type, mT/mG^Tg/+^, and mT/mG^Tg/Tg^ littermates incubated in HBSS containing AnV in the absence (control) or the presence of 100 μM benzamil. Quantification of the AnV (**E**) and mtdTomato (**F**) fluorescence intensity as in D. In B, C, E and F, mean fluorescence intensity ± SD is represented for n = 126-57 OHCs and 43-21 IHCs from 2 cochleae. One-way ANOVA analysis was performed (n.s. p>0.05, \**p*<0.05 and \*\*\*\**p*<0.0001). Scale bar is 10 μm. See also Figure S4.

### TMC1, but not TMC2, is essential for PS externalization and membrane blebbing in auditory hair cells

Since both TMC1 and TMC2 form functional MET channels (Pan et al., 2013, Kawashima et al., 2011, Jia et al., 2019), we next evaluated the individual role of TMC1 and TMC2 in PS externalization and membrane blebbing. For this purpose, we first analyzed PS externalization in hair cells from TMC1^-/-^ or TMC2^-/-^ mice in an mT/mG^Tg/+^ (Figure 4A-F) or wild type (Figure S4B-D) background at P6, when both TMC1 and TMC2 contribute to functional MET (Kawashima et al., 2011). In the absence of MET blockers (control), AnV staining was not detected in TMC2^-/-^ or TMC1^-/-^ hair cells (Figure 4A-B and 4D-E, and Figure S4B-D) indicating that the absence of TMC1 or TMC2 does not trigger PS externalization. However, like in wild type hair cells, treatment with neomycin, benzamil and curare induced PS externalization and an increase of the mtdTomato signal in hair cells from TMC2^-/-^ mice (Figure 4A-C and Figure S4C-D). In contrast, hair cells from TMC1^-/-^ mice failed to externalize PS after MET channel blockade (Figure 4D-E and Figure S4B-D). Furthermore, MET blockade by benzamil or curare did not increase mtdTomato signal in mT/mG^Tg/+^ TMC1^-/-^ hair cells, whereas neomycin was still able to induce a detectable increase of mtdTomato fluorescence intensity in these cells (Figure 4F). Importantly, this neomycin-induced increase of mtdTomato signal was not observed in hair cells from mT/mG^Tg/+^ TMC1^-/-^ TMC2^-/-^ mice (Figure 3C). Besides its MET-blocking activity, neomycin is known to permeate through functional MET channels and accumulate inside the hair cell causing cell death (Alharazneh et al., 2011). Therefore, permeation of neomycin through TMC2 MET channels could explain these results. Together, these data indicate that TMC1 is required for PS externalization and membrane blebbing at the apical hair cell region whereas TMC2 is dispensable for this effect. To our knowledge, this is the first functional difference identified between TMC1 and TMC2.

**FIGURE 4.**
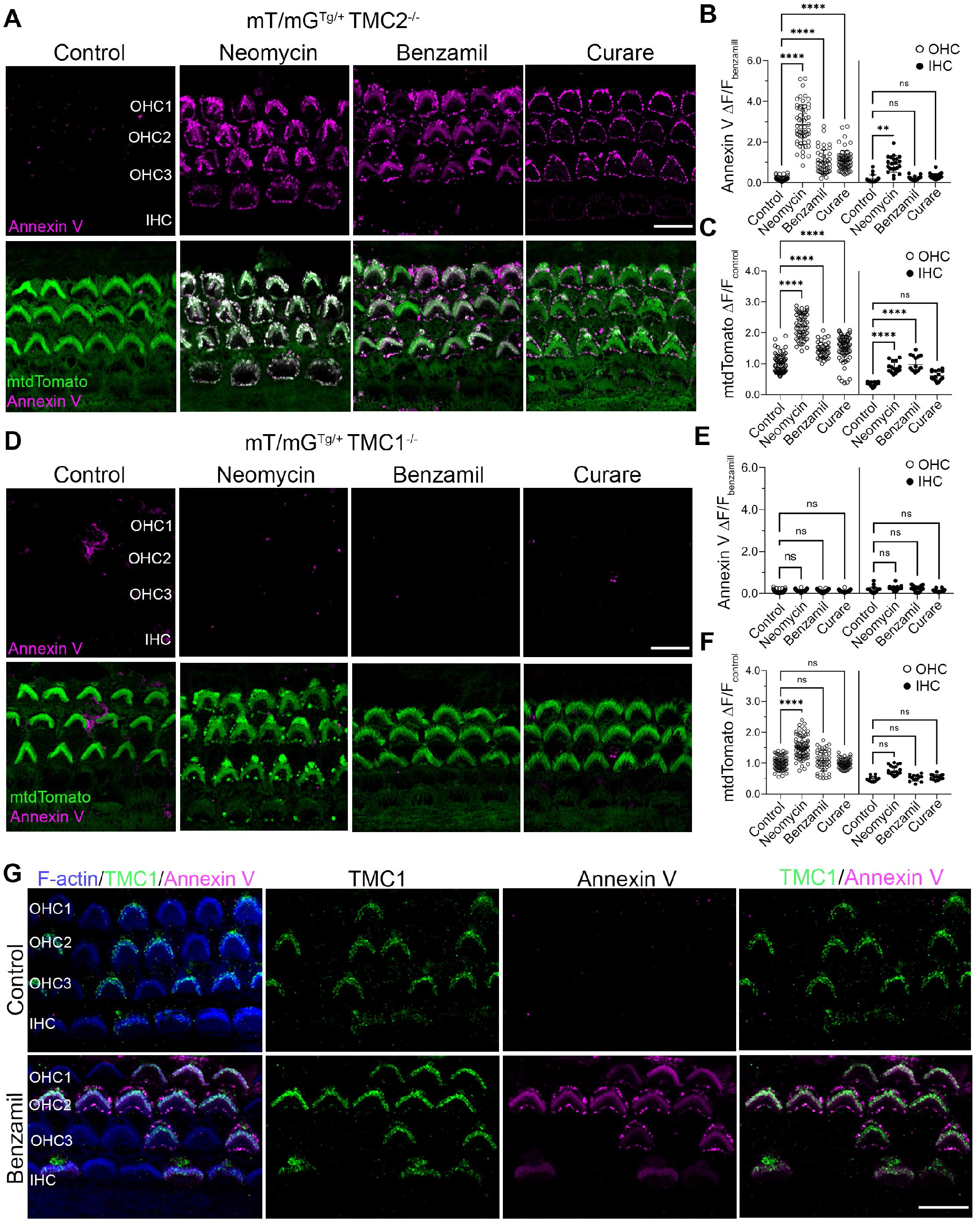
TMC1, but not TMC2, is required for PS externalization and membrane blebbing. **A)** Confocal images of P6 mT/mG^Tg/+^ TMC2^-/-^ hair cells incubated with HBSS containing AnV (magenta) in the absence (control) or the presence of 100 μM neomycin, benzamil or curare. Quantification of AnV (**B**) and mtdTomato (**C**) fluorescence intensity at the apical region of OHCs (○) and IHCs (●) treated as in A. **D)** Confocal images of P6 mT/mG^Tg/+^ TMC1^-/-^ hair cells treated as in A. Quantification of AnV (**E**) and mtdTomato (**F**) fluorescence in the mT/mG^Tg/+^ TMC1^-/-^ mice as in B-C. **G**) Confocal images of hair cells from P6 mosaic mice expressing TMC1-Cherry incubated untreated (control) or treated with 100 μM benzamil. F-actin (blue) is included to visualize the stereocilia. In B, C, E and F, mean ±SD is shown for n =81-45 OHCs and 24-17 IHCs from 2 cochleae. One-way ANOVA analysis was performed (n.s. p>0.05, \*\**p*<0.01 and \*\*\*\**p*<0.0001). Scale bar is 10 μm. See also Figure S4.

After birth, TMC1 expression in OHCs initiates in the basal portion of the cochlea and progresses toward the apex over the subsequent weeks, following the developmental acquisition of MET and the tonotopic organization of the cochlea (Beurg et al., 2018). Consistent with this TMC1-expression gradient of the developing cochlea and with a role of TMC1 in PS externalization, we observed that curare-induced PS externalization in OHCs was stronger in the basal than in the middle cochlear regions whereas it was undetectable in the apical region of the cochlea from P3 wild type mice (Figure S5A-C), when the tonotopic MET gradient differences are still evident (Lelli et al., 2009). Similarly, at this age, PS externalization was only detectable in IHCs from the basal region (Figure S5B-C). Therefore, PS externalization follows the tonotopic gradient of the cochlea. Importantly, we confirmed that PS externalization was not restricted to the immature auditory system since hair cells from P14 mT/mG^Tg/+^ mice also externalized PS in response to the MET blocker benzamil (Figure S5D).

Finally, to further confirm the specific role of TMC1 in PS externalization and membrane remodeling, we analyzed the effect of benzamil on the membrane of hair cells from transgenic mice with mosaic expression of the TMC1-cherry transgene, which does not affect MET (Kurima et al., 2015). In these mice, all hair cells are TMC1^-/-^ TMC2^+/+^ and only some of them express TMC1 fluorescently tagged with a cherry fluorescent reporter (Makabe et al., 2020). Using an anti-cherry antibody, we identified the TMC1-cherry-expressing cells and located the expression of this transgene at the tips of the two shorter stereocilia rows of OHCs and IHCs (Figure 4G), consistent with the well-known localization of TMC1 (Kurima et al., 2015). We did not detect AnV staining in HBSS-treated cells (control), indicating that expression of the TMC1-cherry transgene does not alter PS distribution (Figure 4G). In contrast, benzamil treatment triggered PS externalization only in TMC1-cherry-expressing hair cells, confirming the unique role of TMC1 in the regulation of hair cell membrane homeostasis.

### Buffering of intracellular Ca^2+^ triggers PS externalization and membrane blebbing

PS externalization by TMEM16 scramblases is triggered by a rapid rise in intracellular Ca^2+^ concentration (Suzuki et al., 2010, Yang et al., 2012). In contrast, we have demonstrated that PS externalization in hair cells is induced after MET blockade or breakage of the tip link, treatments that diminish Ca^2+^ influx through the MET channels and thus exclude a role of Ca^2+^ activated TMEM16 scramblases in the process studied here. To determine whether a decrease in intracellular Ca^2+^ plays a role in PS externalization, we incubated hair cells from wild type (Figure S3D-E) and mT/mG^Tg/+^ mice (Figure 5) with the membrane-permeable Ca^2+^ chelator, BAPTA-AM, which does not affect the integrity of the tip links or MET (Velez-Ortega et al., 2017, Peng et al., 2013). As a control, we incubated explants with an equivalent amount of the vehicle used to solubilize BAPTA-AM, but as shown in Figures 5 and S3D, no AnV staining was detectable in these vehicle-treated control hair cells. In contrast, BAPTA-AM induced robust PS externalization at the apical hair cell region and an increase in mtdTomato fluorescence intensity (Figure 5), indicating that buffering of intracellular Ca^2+^ triggers PS externalization and membrane blebbing. However, we found that, like the MET blockers, BAPTA-AM failed to induce PS externalization or an increase of the mtdTomato signal in hair cells from mT/mG^Tg/-^ TMC1^-/-^ mice (Figure 5B-D). Thus, we conclude that externalization triggered by MET blockade or intracellular Ca^2+^ buffering shares a common mechanism that requires TMC1.

**FIGURE 5.**
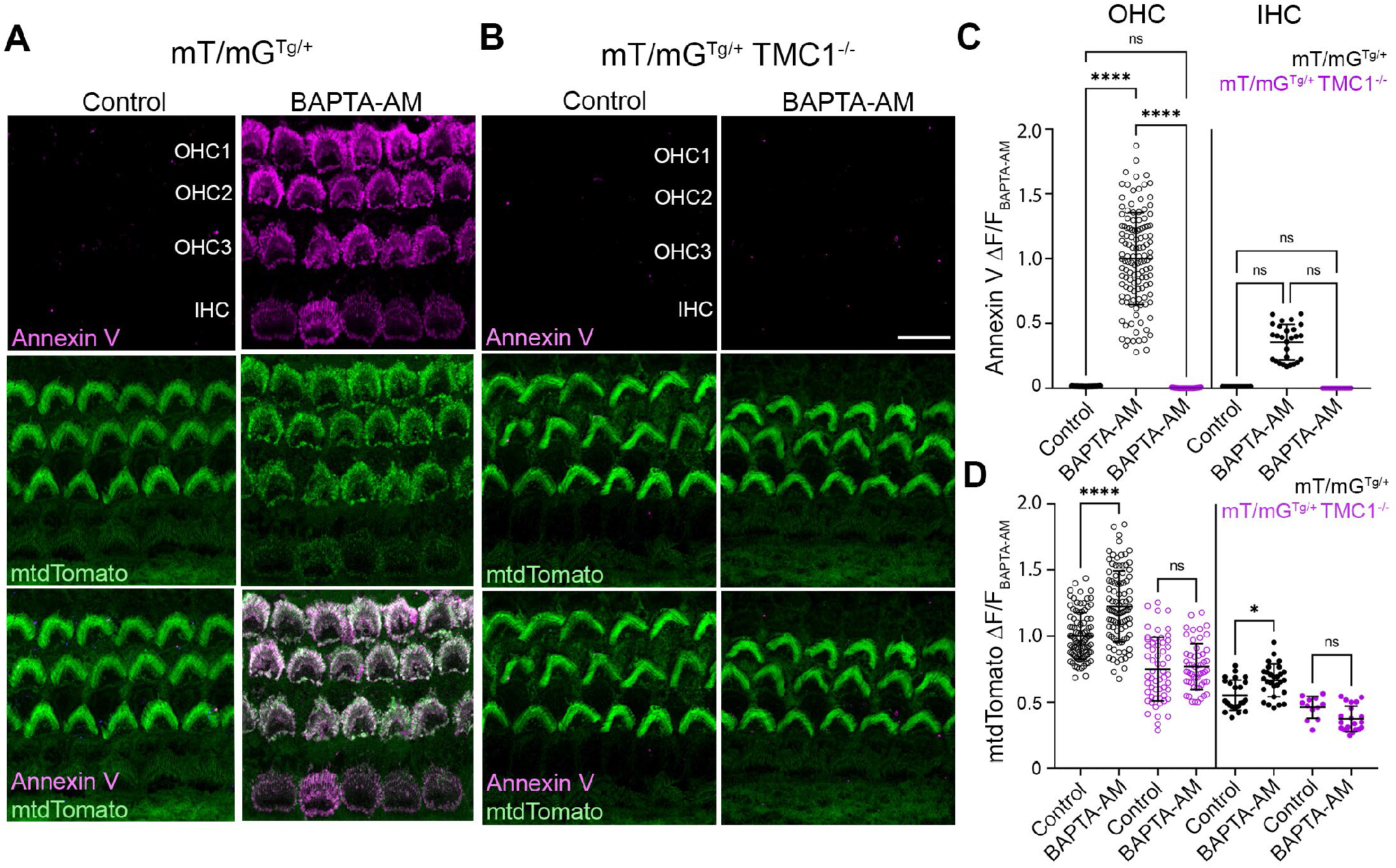
Buffering of intracellular Ca^2+^ triggers PS externalization and membrane blebbing when TMC1 is present. Confocal images of P6 mT/mG^Tg/+^ (**A**) or mT/mG^Tg/+^ TMC1^-/-^ (**B**) hair cells incubated with HBSS containing AnV (magenta) in the presence of BAPTA-AM or the vehicle (Control). Scale bar is 10 μm. Quantification of AnV (**C**) and mtdTomato (**D**) fluorescence intensity at the apical region of OHCs (○) and IHCs (●) treated as in A-B. Mean fluorescence intensity ± SD is represented for n = 70-100 OHCs and 15-30 IHCs from 2 cochleae. One-way ANOVA analysis was performed (n.s. p>0.05, \**p*<0.05, and \*\*\*\**p*<0.0001). See also Figure S3.

### TMC1 deafness-causing mutations cause constitutive PS externalization

Next, we investigated the effect on this TMC1-dependent hair cell membrane remodeling of three TMC1 single-point mutations known to cause deafness in human and mice with different penetrance. We analyzed PS externalization in hair cells from mice carrying the dominant M412K (also termed Beethoven) or D569K mutations, or the recessive D528N mutation in homozygosity and heterozygosity. To compare the degree of PS externalization on all TMC1 genotypes, the AnV signal of untreated hair cells was normalized with the signal of benzamil-treated hair cells for each genotype. Surprisingly, we found that, unlike wild type hair cells, cells homozygous for any of these 3 mutations presented high levels of externalized PS even in the absence of benzamil (control) (Figure 6), indicating constitutive PS externalization at the apical hair cell region of these mutants. More strikingly, this constitutive PS externalization was also detected in untreated hair cells from mice heterozygous for the two dominant TMC1 mutations (TMC1^M412K/+^ and TMC1^D569N/+^) (Figure 6A and 6B). In contrast, like wild type cells, hair cells carrying just one copy of the recessive D528N mutation (TMC1^D528N/+^) lacked externalized PS and required MET blockade with benzamil to externalize PS (Figure 6C). Of note, consistent with our previous results showing that TMC2 is dispensable for PS externalization, we also confirmed that differences in the TMC2 genotype did not affect constitutive PS externalization in TMC1^M412K/+^ hair cells (Figure 6D-E and Figure S6A-B). Importantly, in the absence of benzamil, no significant differences in the AnV signal were found between the homozygous and heterozygous genotypes of the dominant TMC1 mutations, whereas AnV staining was only detectable on hair cells from mice homozygous for the recessive mutation D528N (TMC1^D528N/ D528N^)(Figure 6D and 6E). Therefore, the constitutive PS externalization observed at the apical hair cell region in these TMC1 mutants parallels their expression of the deafness phenotype. Furthermore, while benzamil-treatment further increased the AnV signal in TMC1^M412K/+^ and TMC1^M412K/M412K^ hair cells, blockade of MET did not enhance PS externalization in TMC1^D569N/+^, TMC1^D569N/D569N^ or TMC1^D528N/ D528N^ OHCs (Figure 6D), suggesting a more efficient constitutive externalization of PS in these latter genotypes. However, benzamil potentiated PS externalization in IHCs of all genotypes (Figure 6E). Our data demonstrate that these three TMC1 deafness-causing mutations alter membrane homeostasis and activate constitutive PS externalization in direct correlation with their deafness phenotype. We thus propose that PS externalization could be a critical step in the molecular mechanism of hearing loss.

**FIGURE 6.**
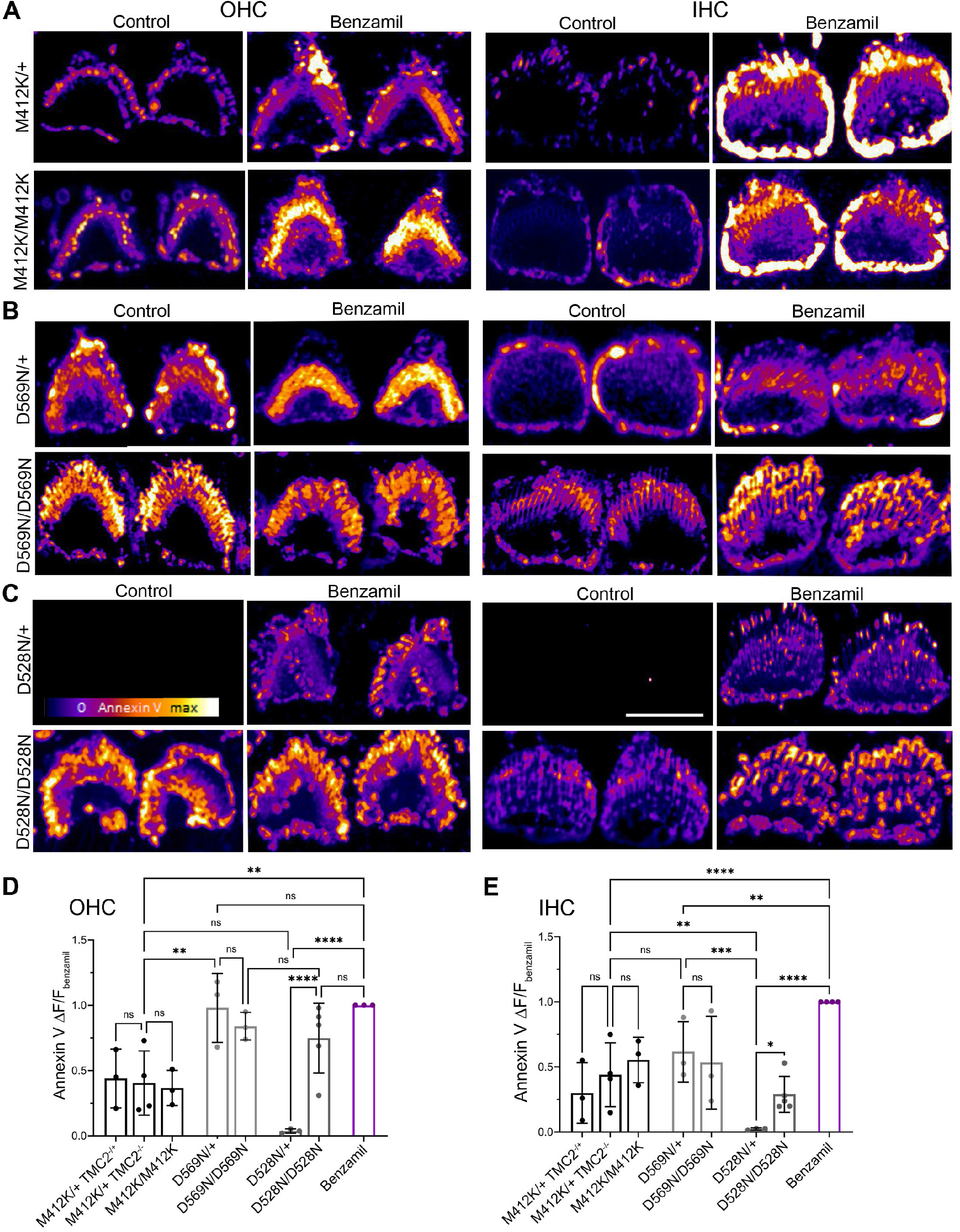
TMC1 deafness-causing mutations render constitutive externalized PS. Confocal images of P6 TMC1^M412K/+^ and TMC1^M412K/M412K^ (**A**), TMC1^D569N/+^ and TMC1^D569N/D569N^ (**B**), or TMC1^D528N/+^ and TMC1^D528N/D528N^ (**C**) outer (OHC) and inner (IHC) hair cells labeled with AnV in the absence (control) or the presence of 100 μM benzamil. Scale bar is 5 μm. Quantification of AnV fluorescence intensity in OHCs (**D**) or IHCs (**E**). Mean ± SD from 3-5 mice is shown. Each dot represents the mean fluorescence intensity of >100 OHCs or >30 OHCs from one cochlea. One-way ANOVA analysis was performed (n.s. p>0.05, \**p*<0.05, \*\**p*<0.01). See also Figure S6.

### PS externalization alters TMC1 localization at the stereocilia tips

To examine how this TMC1-dependent PS externalization, membrane blebbing and ectosome release might cause deafness, we analyzed the TMC1 localization and PS externalization in hair cells over time after MET blockade with benzamil (Figure 7). For these experiments, since immunostaining of endogenous TMC1 has proved to be challenging due to the lack of sensitive and specific antibodies, we used the specific anti-cherry antibody to track the expression of TMC1-cherry in P6 mice expressing the TMC1-cherry transgene in all hair cells (Figure S7D). In the absence of benzamil (control), TMC1 localized to the tips of the two shorter stereocilia rows in OHCs and IHCs, consistent with previous studies (Beurg et al., 2018, Kurima et al., 2015), and no externalized PS was detectable by AnV staining (Figure 7 and S7). However, 10 min after benzamil treatment, we observed externalized PS puncta overlapping with TMC1 at the tips of the shorter stereocilia rows (Figure 7A-B and S7), suggesting that PS initially externalizes near the MET channel. To confirm that PS externalization does not occur at the tips of the longest stereocilia row, we looked at the localization of ESP8, a protein known to be enriched at the longest stereocilia (Furness et al., 2013). As shown in Figure S7A, ESP8 puncta at the tips of the longest stereocilia did not overlap with AnV labeling in P6 IHCs. Furthermore, we found a significant direct correlation between the AnV and TMC1 fluorescence intensities (Figure S7B), suggesting that the expression level of TMC1 determines the extent of PS externalization. These data support that TMC1 is intimately involved in the process of PS externalization.

**FIGURE 7.**
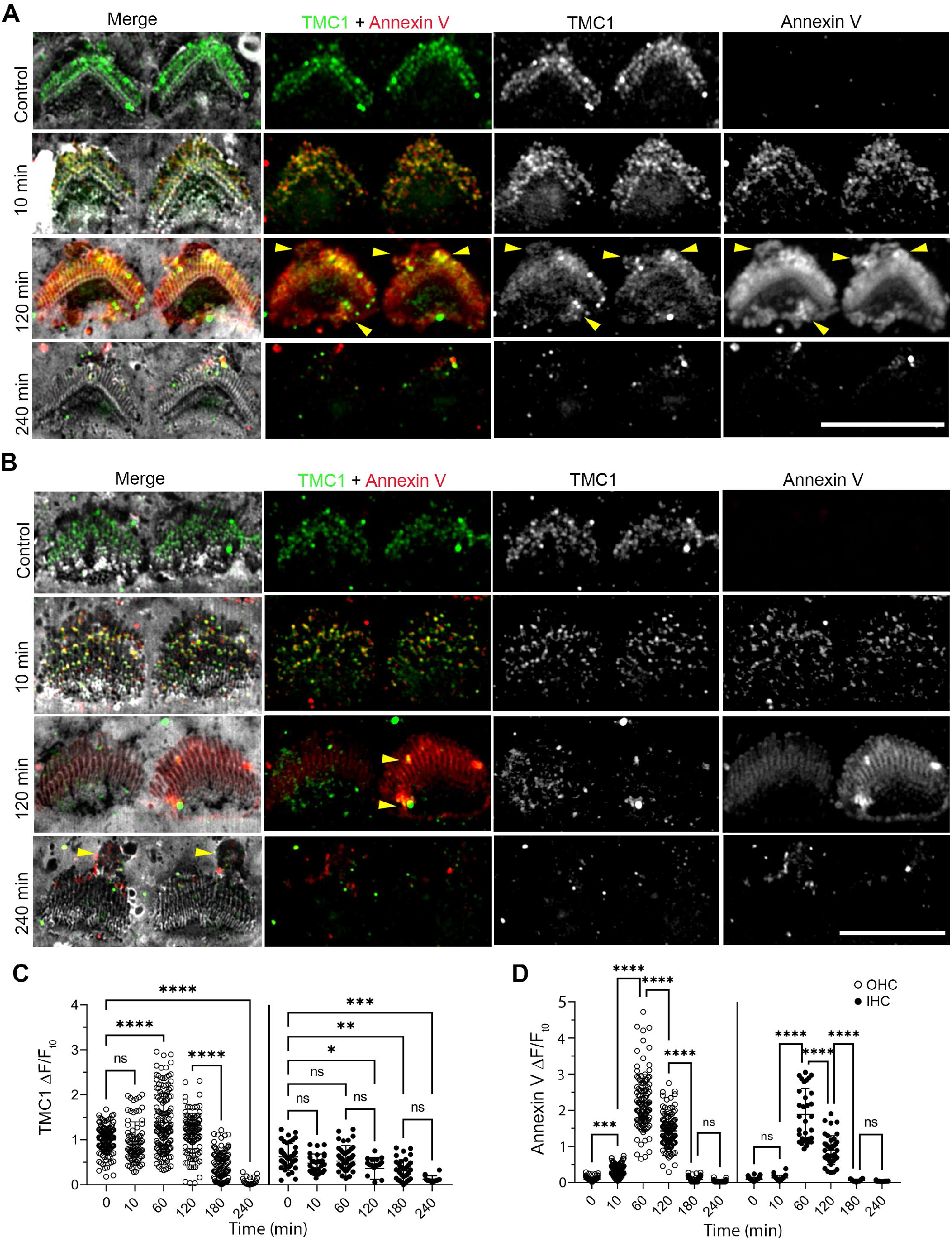
MET blockage and PS externalization alter TMC1 localization at the stereocilia tips. Confocal images of OHCs (**A**) and IHCs (**B**) from P6 TMC1-cherry mice untreated (control) or treated with 100 μM benzamil for 10, 120, and 240 min. TMC1-cherry (green), AnV (red), and WGA (grey) channels are shown. Scale bar is 5 μm. Yellow arrows indicate protruding TMC1 and AnV positive vesicles. Quantification of TMC1 (**C**) and AnV (**D**) fluorescence intensity at the apical region of OHCs (○) and IHCs (●) treated as in A-B. Mean ±SD is shown for n = 125-79 OHCs and 45-23 IHCs from 2 cochleae. One-way ANOVA analysis was performed (n.s. p>0.05, \**p*<0.05, \*\**p*<0.01, \*\*\**p*<0.001, and \*\*\*\**p*<0.0001). See also Figure S7.

Later after benzamil treatment, the PS and TMC1 localization patterns changed. After 1-2 h, AnV labeling and TMC1 remained at the stereocilia but TMC1 lost its puncta-like pattern and it appeared enriched in vesicle-like AnV-positive particles at the stereocilia tips and around the cuticular plate (Figure 7 and S7). After 3-4 h, PS and TMC1 were no longer found at the stereocilia and instead large AnV- and TMC1-positive vesicle-like particles appeared at the kinocilium site and around the cuticular plate (Figure 7 and S7). These results suggest that hair cells release TMC1 in PS-positive ectosomes upon activation of PS externalization by MET blockade. Consistent with this, we found that the signal of WGA staining of the apical hair cell region declined over time after benzamil treatment, indicating loss of glycosylated plasma membrane proteins probably due to ectosome release (Figure S7C). 4h after MET blockade, WGA labeling revealed bulky vesicles at the back of the stereocilia resembling those previously observed by scanning electron microscopy after treatment with ototoxic aminoglycosides (Goodyear et al., 2008, Richardson and Russell, 1991). These data indicate that blockade of the MET channel displaces TMC1 from the stereocilia tips, the cellular localization where TMC1 is required for MET and hearing, by a mechanism engaged following PS externalization.

## DISCUSSION

Our study identifies a novel role of TMC1 MET channels in the regulation of membrane homeostasis in mammalian auditory hair cells. Our data reveal a TMC1-dependent and TMC2-independent mechanism activated by a decrease in intracellular Ca^2+^ that triggers membrane remodeling, and lipid and protein mislocalization. Furthermore, we show that the TMC1 M412K, D569N and D528N deafness-causing mutations alter the membrane regulatory activity of TMC1 according to their dominant or recessive phenotype, suggesting that this new activity of TMC1 is intimately connected with the mechanisms of deafness. These findings provide an integrated view of how hair cell membrane homeostasis is regulated and have important implications for the role of TMC1 in hair cell physiology and pathology.

MET is essential for stereocilia remodeling and proper localization of protein at specific stereocilia rows during bundle development. TMC2 or TMC1 MET channels can support bundle development at early stages (Beurg et al., 2018) but stereocilia dimensions and distribution of stereocilia proteins are refined until mature MET is reached, coinciding with the protein switch from TMC2 to TMC1 (Krey et al., 2020). Since TMC1 channels exhibit lower Ca^2+^ permeability and conductance than TMC2 (Kim and Fettiplace, 2013; Pan et al., 2013), we propose that TMC1 expression might trigger small-scale membrane remodeling and ectosome release to redistribute or remove specific stereocilia proteins for the establishment of the mature sensory apparatus. Consistent with this hypothesis, the transducing stereocilia tips of TMC1^-/-^ hair cells display accumulation of TMC2 and altered protein distribution (Beurg et al., 2018; Krey et al., 2020), supporting a role for TMC1 in the removal of TMC2 and other proteins from the stereocilia. In addition, although we did not detect membrane blebbing in control untreated hair cells, small vesicles have been observed by scanning electron microscopy at the stereocilia of developing hair cells (Velez-Ortega et al., 2017, Waguespack et al., 2007). Interestingly, while the independent role of TMC1 and TMC2 in regulating stereocilia dimensions is unexplored, the immature bundle morphology and altered protein distribution of MET-lacking hair cells can also be achieved by pharmacological blockade of the MET channel, suggesting a common mechanism for bundle maturation and stereocilia remodeling (Krey et al., 2020; McGrath et al., 2021; Velez-Ortega et al., 2017). The lipid and protein mislocalization, and the loss of TMC1 that we demonstrate occur after MET blockade could favor the return of an immature morphology and protein distribution in these hair cells. Alternatively, continuous removal of TMC1 or failure to establish the mature MET complex due to prolonged MET blockade or aminoglycoside exposure might lead to hair cell degeneration and death. Overall, our findings suggest that stereocilia might use membrane remodeling and ectosome release as a mechanism to regulate membrane and protein homeostasis, resembling cilia (Wood and Rosenbaum, 2015). Consistent with this, it has been shown that myosin 6 and small GTPases implicated in ectosomes shedding are required for hair bundle morphogenesis and hearing (Kollmar, 1999; Meldolesi, 2018). Further investigation will be required to characterize the molecular players responsible for membrane remodeling and ectosome release in hair cells and their implications in stereocilia development and repair.

The characterization of TMC1 and TMC2 single and double knockout mice revealed that TMC2 can compensate for the lack of TMC1 in the vestibular but not in the auditory system (Kawashima et al., 2011). Initially, it was proposed that this was due to the loss of TMC2 expression in mature auditory hair cells, but it has been recently shown that transgenic expression of TMC2 in TMC1^-/-^ mice does not rescue auditory function (Nakanishi et al., 2018, Zheng and Holt, 2021). Therefore, TMC1 must have additional functions absent in TMC2 essential for hearing. Here we identify the first functional difference between TMC1 and TMC2 that could explain the different requirement for these two proteins in proper hearing or balance. Unlike vestibular hair cells, auditory hair cells are tonotopically organized along the cochlea and tuned to sense and amplify precise frequencies. Interestingly, we demonstrate that TMC1-dependent PS externalization follows the tonotopic gradient of the cochlea. While our study focused on PS externalization, loss of the membrane asymmetry causes changes in membrane thickness and externalization of other phospholipids such as phosphatidylinositols, which are known to regulate MET adaptation (Gianoli et al., 2017, Hirono et al., 2004, Effertz et al., 2017, Caprara et al., 2020). The ability of TMC1 to remodel the membrane and alter the lipid distribution may equip TMC1 channels to respond and adapt to different frequencies. Accordingly, TMC1 channels show better adaptation than TMC2, and TMC1 deafness-causing mutations affect MET adaptation (Goldring et al., 2019). Although our work reveals that TMC1 MET channels alter membrane homeostasis in a timescale of minutes, it will motivate future efforts to understand the implications for hearing of the relationship between the MET channels and the hair cell membrane at shorter times.

We demonstrate here for the first time that TMC1 deafness-causing mutations present constitutive PS externalization, linking this novel TMC1 activity with the mechanisms of deafness and suggesting that maintenance of hair cell membrane homeostasis is essential for hearing. Consistent with this, it has been shown that mutations in flippases, which transport PS from the outer to inner membrane leaflet, cause hearing loss (Stapelbroek et al., 2009). Auditory hair cells lacking TMC1 or carrying TMC1 deafness-causing mutations degenerate and die leading to hair cell loss, but the molecular mechanisms leading to hair cell degeneration and deafness are unknown. While wild type hair cells might use PS externalization and subsequent membrane blebbing and TMC1 removal to restore malfunctioning MET or prevent aminoglycoside overload and cell death (Kenyon et al., 2021), we propose that constitutive PS externalization in TMC1 M412K, D569N and D528N mutant hair cells may be the underlying mechanism leading to progressive hair cell loss, and ultimately, deafness in these mice and humans carrying the equivalent mutations (M418K and D572N). In fact, reduced TMC1 protein levels and immature-like bundles have been described in TMC1 D569N mice and hair cell loss can be observed in mice homozygous for any of these mutations as early as P15 (Beurg et al., 2019, Beurg et al., 2021, Marcotti et al., 2006). Furthermore, while OHCs are lost before IHCs in TMC1 D569N mice, the opposite has been reported for M412K, agreeing with the levels of PS externalization we observed in the OHCs and IHCs from these mutant mice. In contrast, this correlation between PS externalization and cell loss was not applicable to mice homozygous for the recessive TMC1 D528N mutation, where, although IHCs are lost first, we observed stronger PS externalization in OHCs than in IHCs, suggesting an indirect mechanism for hair cell loss in mice carrying this recessive mutation. Furthermore, our data support that constitutive PS externalization in these dominant and recessive TMC1 mutations is mediated by different mechanisms. In homozygosity, despite its recessive deafness phenotype, D528N alters the MET channel current and Ca^2+^ permeability more severely than the dominant M412K and D569N mutations (Beurg et al., 2021). This suggests that like in wild type hair cells upon MET blockade, PS externalization in D528N homozygous hair cells is caused by a substantial decrease in intracellular Ca^2+^. In contrast, in heterozygosity, despite Ca^2+^ permeability being equally diminished by the recessive D528N or dominant M412K and D569N mutations (Table S1), we observed constitutive PS externalization only in hair cells carrying the dominant mutations, indicating that the changes in MET channel Ca^2+^ permeability are not enough to explain the constitutive PS externalization observed in the dominant mutations. These data support that while recessive D528N is a loss-of-function mutation that results in markedly reduced Ca^2+^ permeability of the MET channel, dominant M412K and D569N are gain-of-function mutations that cause constitutive PS externalization independently of intracellular Ca^2+^. Based on these observations, it is tempting to speculate that TMC1 could moonlight as a MET channel and a Ca^2+^-inhibited lipid scramblase that it is turned into a constitutive scramblase by the dominant mutations. Notably, TMC1 dominant deafness-causing mutations localize to a region that corresponds to the cavity implicated in lipid scrambling in TMEM16 (Ballesteros et al., 2018, Gyobu et al., 2017). Future experiments with purified protein reconstituted in liposomes will help establish the potential scramblase function of TMC proteins more decisively.

Our study reveals new insights into the importance of the hair cell membrane homeostasis for hearing. By studying the interplay between the MET channel and the membrane, we uncovered a TMC1-dependent autoregulatory process that provides a new foundation for elucidating the mechanisms of hair cell bundle development, stereocilia remodeling and deafness-causing mutations.

## ACKNOWLEDGMENTS

We greatly appreciate the feedback and discussions from our lab mates and colleagues, and Helena Chang’s support with the mouse colony. We thank Dr. Andrew Griffith, Dr. Wade Chien, and Dr. Robert Fettiplace for sharing the TMC1 transgenic mice with us. Dr. Leonid Chernomordik kindly provided the Lact-C2 plasmid and purification protocol. Dr. Vincent Schram provided help and feedback with imaging. This study was supported by the Intramural Research Program number NS002945 of the NINDS, NIH, Bethesda, MD, to K.J. S. We apologize to colleagues whose important and relevant work was not cited due to space limitations.

## AUTHOR CONTRIBUTIONS

A.B conceived the research, designed the experiments, analyzed the data and carried out the experiments. K.S contributed to experiments design and interpretation. A.B. and K.S wrote the manuscript.

## MATERIALS AND METHODS

### Mice strains

Wild type C57BL/6J (strain 000664) and mT/mG mice (strain 007676) were purchased from The Jackson Laboratory and breed in our animal facility to obtain the desired phenotypes. TMC1^-/-^, TMC2^-/-^ (RRIDs: IMSR_JAX:019146 and IMSR_JAX:019147) (Kawashima et al., 2011), and transgenic mice endogenously expressing TMC1 fused at the C-terminal to Cherry fluorescent protein in a TMC1^-/-^ TMC2^-/-^ background (TMC1-cherry, RRID: IMSR_JAX:028392)(Kurima et al., 2015) were obtained from Dr. Andrew Griffith (NIDCD). TMC1^-/-^ TMC2^-/-^ mice were crossbreed with mT/mG^Tg/Tg^ mice to obtain double (mT/mG^Tg/Tg^ TMC1^-/-^ and mT/mG^Tg/TG^ TMC2^-/-^) and triple (mT/mG^Tg/Tg^ TMC1^-/-^ TMC2^-/-^) transgenic mice. Since only heterozygous mice for the mtdTomato transgene (mT/mG^Tg/+^) were used in most of our experiments except as indicated, mT/mG^Tg/Tg^ and TMC1^-/-^ mice were crossbred with TMC1^-/-^ to obtain heterozygous mT/mG^Tg/+^ TMC1^-/-^ mice. The same protocol was followed to generate mT/mG^Tg/+^ TMC2^-/-^ and mT/mG^Tg/+^ TMC1^-/-^ TMC2^-/-^ mice. TMC1^M412K/M412K^, TMC1^D569N/D569N^ and TMC1 ^D528N/D528N^ mice were bred with wild type to obtain heterozygous mice. For all genotypes, a mixture of male and female mice was used. Mice were kept on a 12-hour light/dark cycle and were allowed solid food and water ad libitum. The animal care and experimental procedures were performed following the Guide for the Care and Use of Laboratory Animals. They were approved by the Animal Care and Use Committee of the National Institute of Neurological Disorders and Stroke (Animal protocol number 1336).

Genomic DNA extraction from tail snips and genotyping PCR reactions were performed using MyTaq Extract-PCR kit (Bioline, Taunton, MA). All the mice were genotyped by the fragment PCR method as indicated in the Jackson Laboratory website, previously described (Ballesteros et al., 2018), or using the primers listed below. PCR products were run on 2% agarose gels and the Quick load 100pb DNA ladder (New England Biolabs Inc., Ipswich, MA) was used for fragment size visualization. The PCR fragments obtained from genotyping of the TMC1 M412K mice were purified and sent for sequencing, whereas the PCR fragments from the TMC1 D569N or D528N mice were further digested with EcoRI or Acl1, respectively.

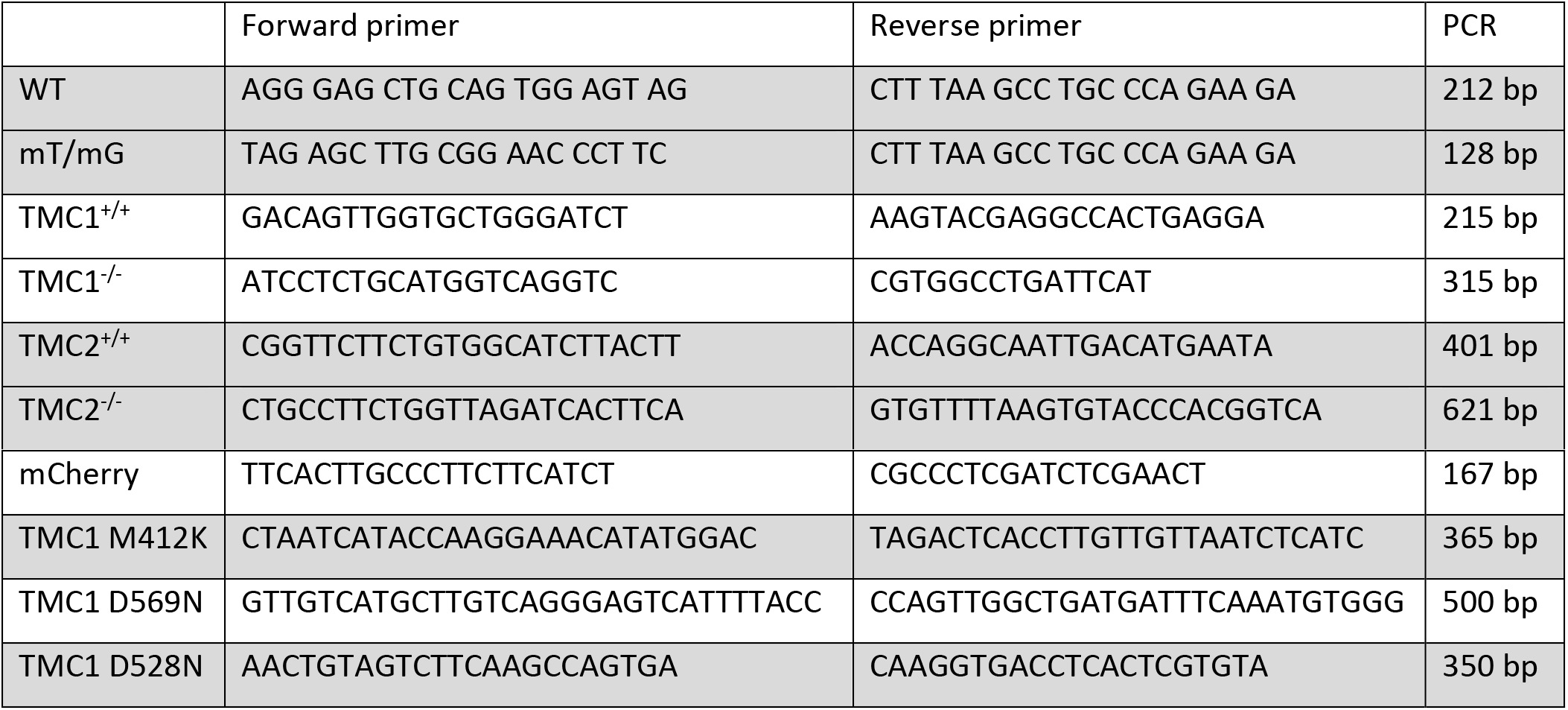

### Dissection of inner ear tissue and sample preparation for imaging

Excision of the inner ear from P6 wild type C57BL/6J, reporter mT/mG, or TMC mutant mice and further cochleae dissection, including removing the semicircular canals and vestibular organs were performed in cold Leibovitz’s L15 media. Cochlear tissues were placed on a corning PYREX 9 depression plate well Hanks’ Balanced Salt solution (HBSS) to remove the cochlear bone and expose the organ of Corti. At least three cochleae from littermate mice were used for each experimental condition. Drug treatments were performed at this stage. The cochlear tissues were incubated with the indicated drugs and fluorescent dyes for 25 min at RT in an orbital shaker and protected from the light. After treatment, tissues were washed twice with HBSS, and fixed in 4 % paraformaldehyde (PFA) in HBSS for 30 min. Fixed tissue was washed with HBSS to remove PFA and the spiral ligament and the tectorial membrane were removed to obtain fixed organ of Corti explants in HBSS buffer. Tissue was washed with Phosphate-Buffered Saline (PBS) to remove salt and finally mounted with ProLong Diamond antifade mounting media (ThermoFisher Scientific) on a superfrost plus microscope slide (Fisherbrand) and covered with a #1.5 glass coverslips of 0.17 ± 0.02 mm thickness (Warner Instruments) for confocal imaging.

### Blockage of the MET channel

Dissected organ of Corti from wild type C57BL/6J, mT/mG^Tg/+^ or TMC1 or TMC2 single and double knock out littermates mice were incubated in HBSS containing 1/25 annexin V-Alexa647 (Thermofisher) and 20 μg/mL CF488A-WGA (Biotium) in the absence (control) or presence of neomycin, benzamil, or curare at 0.1 mM concentration. A stock solution of 30 mM benzamil hydrochloride hydrate (Sigma-Aldrich) was prepared in DMSO. Neomycin trisulfate salt hydrate (Sigma-Aldrich) was dissolved at 50 mM in water. Curare or tubocurarine hydrochloride pentahydrate (Sigma-Aldrich) was dissolved at 50 mM in water. Stock solutions were aliquoted and stored at −30 °C. Annexin V was used to detect and visualize the externalized PS due to its ability to bind PS with high affinity in a Ca^2+^-dependent manner (Meers and Mealy, 1993). Wheat germ agglutinin (WGA) binds to oligosaccharides and has been shown to label the surface of both IHCs and OHCs without blocking transduction (Bhavanandan and Katlic, 1979; Richardson et al., 1989). In all the conditions we tested, WGA labeled the stereocilia and hair cell surface and was also accumulated at the surface of pillar and Hensen cells (Figure 1A). P6 heterozygous mT/mG mice (mT/mGTg/+), which express a membrane-targeted tandem dimer tomato (mtdTomato) fluorescent protein and preserve MET (Ballesteros et al., 2021) were used to visualize the stereocilia and hair cell membrane. Cochlear tissues were incubated with the drugs and fluorescent dyes for 25 min at RT in an orbital shaker protected from light. After treatment, tissues were washed twice with HBSS, fixed in 4 % PFA in HBSS for 30 min, and prepared for imaging as described above. P6 littermates mice obtained from mT/mG^Tg/+^ and mT/mG^Tg/+^ breedings were used in the experiments comparing wild type, mT/mG^Tg/+^ and mT/mG^Tg/Tg^ hair cells. One cochlea was incubated in HBSS, and the other one was treated with benzamil. Cochleae from wild type or TMC2^-/-^ mice were included in the experiments with TMC1^-/-^ and TMC1^-/-^ TMC2^-/-^ mice as positive controls.

For live imaging experiments, dissected organs of Corti from P4-P6 wild type mice were cultured over collagen covered 35 mm glass-bottom cell culture dish with 2 mL of Leibovitz’s L15 media with fetal bovine serum (FBS) at 37 °C and 5% CO_2_. After 2-4h incubation, 5 μL of CellMask Green plasma membrane stain (ThermoFisher) was added to the Leibovitz’s L15 media with FBS and incubated at 37 °C for 5-10 min. Cell media was removed, and 1/50 annexin V was added in Leibovitz’s L15 media without FBS. After collecting some images of time cero, 100 μM curare was added to the annexin V containing media, and imaging acquisition was resumed at RT. An equivalent amount of water was added to the control samples.

### Disruption of the tip links with BAPTA

We incubated organ of Corti explants with 5 mM BAPTA in HBSS without Ca^2+^ and Mg^2+^ (HBSS-CFM) containing CF488A-WGA at 20 μg/mL for 25 min. A stock of BAPTA tetrapotassium salt (Molecular Probes, ThermoFisher scientific, Inc.) was dissolved at 50 mM in water, aliquoted, and stored at −30 °C. Organ of Corti explants were incubated in HBSS or HBSS-CFM buffer containing CF488A-WGA at 20 μg/mL as controls. Tissue explants were then washed once with HBSS, and annexin V-Alexa647 was added at a 1/25 dilution in HBSS for 5 min. Tissues were washed twice with HBSS to remove the excess of annexin V, fixed in 4 % PFA in HBSS for 30 min, and prepared for imaging as described above. In addition, experiments with the C2 domain of lactadherin fused to a fluorescent clover protein (clover-Lact-C2) were performed to detect the externalized PS in the absence of Ca^2+^. In these experiments, excised organs of Corti tissues were incubated in HBSS-CFM with 20 μg/mL CF488A-WGA and 3 μg/mL clover-Lact-C2 for 25 min at RT under gentle shaking. Tissues were then washed twice with HBSS to remove the excess of WGA and clover-Lact-C2, fixed in 4 % PFA in HBSS for 20 min, and prepared for imaging as described above.

### Expression and purification of clover-Lact-C2

A cDNA construct consisting of Clover fluorescent protein followed by a hexa-histidine tag and lactadherin-C2 (Clover-(His)6-LactC2) in the pET-28 bacterial expression vector together with pLysSRARE2 plasmid (both kindly shared by Dr. Chernomordik, NICHD) were electroporated into ClearColi™ BL21(DE3) according to manufacturer protocol (Lucigen, WI). Bacterial colonies that grew in an LB plate containing 100 μg/ml kanamycin and 30 μg/ml chloramphenicol, were considered positive for incorporating both plasmids and selected for protein expression. Cells were grown in 0.5liter of LB Miller medium at 37°C in 100 μg/ml kanamycin and 30 μg/ml chloramphenicol until the culture reached A600 = 1. After the addition of 1 mM IPTG, the culture was grown for 3 hr at 37°C. Next, cells were lysed in B-PER bacterial protein extraction reagent (Thermo Scientific, MA) containing 2500 U of DNAseI (Sigma), 0.1 mM of PMSF, and SIGMAFAST EDTA-free protease inhibitor cocktail (Sigma). After centrifugation at 20,000 × g for 20 min, Clover-(His)6-LactC2 was purified from the supernatant with 3mL of the TALON superflow metal affinity resin (Clontech, CA). The resin was washed with 5 mM imidazole in PBS, and the protein was eluted with 100 mM of imidazole in PBS (Figure S3C). The eluted protein was dialyzed against PBS and concentrated to a final concentration of 1.036 mg/mL.

### Perilymph and endolymph-like buffers

The perilymph-like buffer used in our experiments had a composition of 125 mM NaCl, 1.3 mM CaCl_2_, 3.5 mM KCl, 1.5 mM MgCl_2_, 10 mM HEPES pH 7.4, 5 mM Glucose, and the osmolarity was adjusted to 290 mOs with NaCl. The composition of the endolymph-like buffer used was: 1 mM NaCl, 0.025 mM CaCl_2_, 126 mM KCl, 0.025 mM MgCl_2_, 10 mM HEPES pH 7.4, 0.5 mM Glucose, and the osmolarity was adjusted to 315 mOs with KCl. Cochlear tissues from P6 mT/mG^Tg/+^ mice were dissected to remove the cochlear bone and expose the organ of Corti. Organ of Corti explants were incubated in perilymph-like or endolymph-like buffer with benzamil or the equivalent volume of DMSO (control) for 25 min at RT with gentle shaking in an orbital shaker protected from the light. Tissue explants were then washed once with HBSS, and annexin V-Alexa647 was added at a 1/25 dilution in HBSS for 5 min to all the samples. Tissues were washed twice with HBSS to remove the excess of annexin V, fixed in 4 % PFA in HBSS for 30 min, and prepared for imaging as described above.

### Buffering of intracellular Ca^2+^

To buffer the intracellular Ca^2+^ concentration in murine hair cells we use the Ca^2+^ quelator BAPTA-AM. BAPTA-AM is an acetoxymethyl (AM) ester derivative of BAPTA that binds Ca^2+^ only after its AM group is removed by cytoplasmic esterases. Importantly, treatment of cochlear tissue with BAPTA-AM does not affect the integrity of the tip links or does not alter the resting probability of the MET channel (Velez-Ortega et al., 2017, Peng et al., 2013). A stock of cell-permeant BAPTA-AM (Invitrogen) was prepared at 20 mM in 20% pluronic F-127 in DMSO, aliquoted, and stored at −30 °C. Pluronic F-127 is a mild non-ionic detergent useful for solubilizing BAPTA-AM and facilitating AM esters loading into cells. A stock solution of HBSS containing 20 μg/mL of CF488A-WGA, and annexin V at 1/25 dilution was prepared and split into two tubes. 20 mM BAPTA-AM in 20% pluronic F-127 in DMSO was added at a final concentration of 20 μM while an equivalent volume of the vehicle (20% pluronic F-127 in DMSO) was added to the other tube and used as a control in these experiments. Dissected organ of Corti explants were incubated with the solution containing BAPTA-AM or the vehicle and incubated for 25 min at RT under gentle shaking. After the incubation, the tissue was washed a couple of times with HBSS, fixed with 4% PFA, and prepared for imaging.

### TMC1-cherry immunostaining and time course

Organ of Corti explants from TMC1-cherry mice were dissected as described above. Tissue explants were then incubated with HBSS (control) or HBSS with 100 μM benzamil in the presence of 1/25 annexin V-Alexa647 and 20 μg/mL CF488A-WGA. 10, 60, 120, 180 and 240 min after benzamil addition, tissues were washed twice with HBSS, and fixed in 4 % PFA in HBSS for 30 min. Fixed tissues were washed with HBSS to remove PFA and the spiral ligament and the tectorial membrane were removed to obtain fixed organ of Corti explants in HBSS buffer. To label TMC1-cherry, tissues were permeabilized in 0.5% Triton X-100 in PBS for 20 min. Tissues were washed twice with PBS for 5 min to remove the excess of Triton X-100 and then blocked for 1h at RT with blocking buffer (PBS containing 10% goat serum and 5% BSA). The primary antibody anti-RFP (660-401-379, Rockland) was added at 1/100 in blocking buffer and incubated at 4 °C overnight in an orbital shaker with gentle shaking protected from light. The next morning, tissues were washed three times with PBS to remove the excess of primary antibody. Goat anti-rabbit secondary antibody conjugated with Alexa Fluor 594 (R37117, Molecular probes) at the manufacturer’s recommended concentration and 1:100 CF405M phalloidin (Biotium) in blocking buffer were added for 30 min at RT. Tissues were then washed 2-3 times with PBS and prepared for imaging as described above.

### PS externalization in TMC1 mutants

To compare the annexin V fluorescence in the TMC1 mutant mice, we incubated one cochlea in HBSS containing annexin V-Alexa647 at a 1/25 dilution and CF488A-WGA (Biotium) at 20 μg/mL and the other cochlea in the same buffer supplemented with 100 μM benzamil for 25 min at RT in an orbital shaker protected from the light. After treatment, tissues were washed twice with HBSS, fixed in 4 % PFA in HBSS for 30 min, and prepared for imaging as described below.

### Quantification and statistical analysis

Super-resolution imaging was performed in the Neuroscience Light Imaging Facility (NINDS) with a confocal laser scanning microscope Zeiss LSM 880 (Carl Zeiss AG) equipped with a 32 channel Airyscan detector. Images of the whole organ of Corti were taken with a 20X objective (Carl Zeiss). To image the hair cells, we used oil immersion alpha Plan-Apochromat 63X/1.4 Oil Corr M27 objective (Carl Zeiss) and immersol 518F media (ne=1.518 (30°)). A z-stack of images from the stereocilia to the apical half of the hair cell body was collected with identical image acquisition settings, no averaging, and optimal parameters for x, y, and z resolution for each independent experiment. Image acquisition and Airyscan image processing were performed with Zen Black 2.3 SP1 software (Carl Zeiss) using the Airyscan 3D reconstruction algorithm with the automatic default Wiener filter settings. Samples from each experiment, including control and treated or wild type and mutant mice, were imaged on the same day to limit variability.

Microscopy data analysis and quantification were done in ImageJ (Schneider et al., 2012). To measure the fluorescence intensity at the apical region of the hair cells, we generated a region of interest (ROI) around each OHCs and IHCs using the circular tool. The fluorescence intensity of an equivalent ROI in an area outside the hair cells was considered as background and subtracted from the values at the hair cells for each image. Annexin V and mtdTomato fluorescence intensities were quantified in each hair cell. mtdTomato fluorescence intensity was normalized against the mtdTomato fluorescence intensity of the control samples, whereas annexin V fluorescence intensity was normalized against the fluorescence of the treated samples as indicated in the corresponding y-axis. The number of cells and tissues analyzed in each condition and the statistical method used are indicated in the figure legends. GraphPad Prism V.7 software was used to generate the graphs and perform the statistical analysis.

## SUPPLEMENTARY INFORMATION

**Table S1.** Summary of the mice strains used in this study and main results. MET blockers treatment includes neomycin, benzamil, and curare. PD is profound deafness and HL is hearing loss due to elevated ABR thresholds. Permeability of Ca^2+^ versus Cs^+^ (PCa^2+/^Cs^+^)* in WT mice was estimated as ^~^5.5.

| Mouse Strain | Treatment | Externalized PS |  | MET | ~PCa/Cs | Phenotype | References |
| --- | --- | --- | --- | --- | --- | --- | --- |
|  |  | Pre treatment | Post treatment |  |  |  |  |
| WT | MET blockers<br>BAPTA/BAPTA-AM | NO | +++ | Yes | 4.25 | WT | Kawashima, Y. et al. J Clin Invest 2011.<br>Pan, B. et al. Neuron. 2014.<br>Kim, KX. et al. Gen Physiol. 2013. |
| mT/mG<sup>+/+</sup> | benzamil | NO | +++ | Yes | — | WT | Ballesteros, A. et al. Hearing Research. 2021. |
| mT/mG<sup>Tg/+</sup> | MET blockers<br>BAPTA/BAPTA-AM | NO | +++ | Yes | — | HL P60 | Ballesteros, A. et al. Hearing Research. 2021. |
| mT/mT<sup>Tg/Tg</sup> | benzamil | NO | NO | No | — | PD | Ballesteros, A. et al. Hearing Research. 2021. |
| TMC1<sup>-/-</sup> | MET blockers | NO | NO | Yes | 6.5* | PD | Kawashima, Y. et al. J Clin Invest 2011.<br>Pan, B. et al. Neuron. 2014.<br>Kim, KX. et al. Gen Physiol. 2013. |
| TMC2<sup>-/-</sup> | MET blockers | NO | +++ | Yes | 4.5* | WT | Kawashima, Y. et al. J Clin Invest 2011.<br>Pan, B. et al. Neuron. 2014.<br>Kim, KX. et al. Gen Physiol. 2013. |
| TMC1<sup>-/-</sup> TMC2<sup>-/-</sup> | MET blockers | NO | NO | No | 0 | PD | Kawashima, Y. et al. J Clin Invest 2011.<br>Pan, B. et al. Neuron. 2014.<br>Kim, KX. et al. Gen Physiol. 2013. |
| TMC1<sup>-/-</sup><br>mT/mG<sup>Tg/+</sup> | MET blockers<br>BAPTA-AM | NO | NO | Yes | — | — | This study |
| TMC2<sup>-/-</sup><br>mT/mG<sup>Tg/+</sup> | MET blockers | NO | +++ | Yes | — | — | This study |
| TMC1<sup>-/-</sup> TMC2<sup>-/-</sup><br>mT/mG<sup>Tg/+</sup> | MET blockers<br>BAPTA-AM | NO | NO | No | 0 | — | This study |
| TMC1-Cherry | benzamil | NO | +++ | Yes | — | WT | Kurima, K. et al. Cell Reports. 2015. |
| TMC1-Cherry<br>mosaic | benzamil | NO | +++ | Yes | — | HL P60 | Kurima, K. et al. Cell Reports. 2015. |
| TMC1<sup>M412K/+</sup> | benzamil | + | +++ | altered | 2.4 | HL P30 | Beurg, M. et al. PNAS. 2019.<br>Pan, B. et al. Neuron. 2014.<br>Vreugde, S. et al. Nat. Genetics. 2002. |
| TMC1<sup>M412K/M412K</sup> | benzamil | + | +++ | altered | 1.5 | PD P30 | Beurg, M. et al. PNAS. 2019. |
| TMC1<sup>D569N/+</sup> | benzamil | +++ | +++ | altered | 2 | PD P30 | Beurg, M. et al. PNAS. 2019. |
| TMC1<sup>D569N/D569N</sup> | benzamil | +++ | +++ | altered | 1.25 | PD P30 | Beurg, M. et al. PNAS. 2019. |
| TMC1<sup>D528N/+</sup> | benzamil | NO | +++ | altered | 2 | WT P60 | Beurg, M. et al. J. Neurosci. 2021. |
| TMC1<sup>D528N/D528N</sup> | benzamil | +++ | +++ | altered | 0.57 | PD P28 | Beurg, M. et al. J. Neurosci. 2021. |

**Figure S1.**
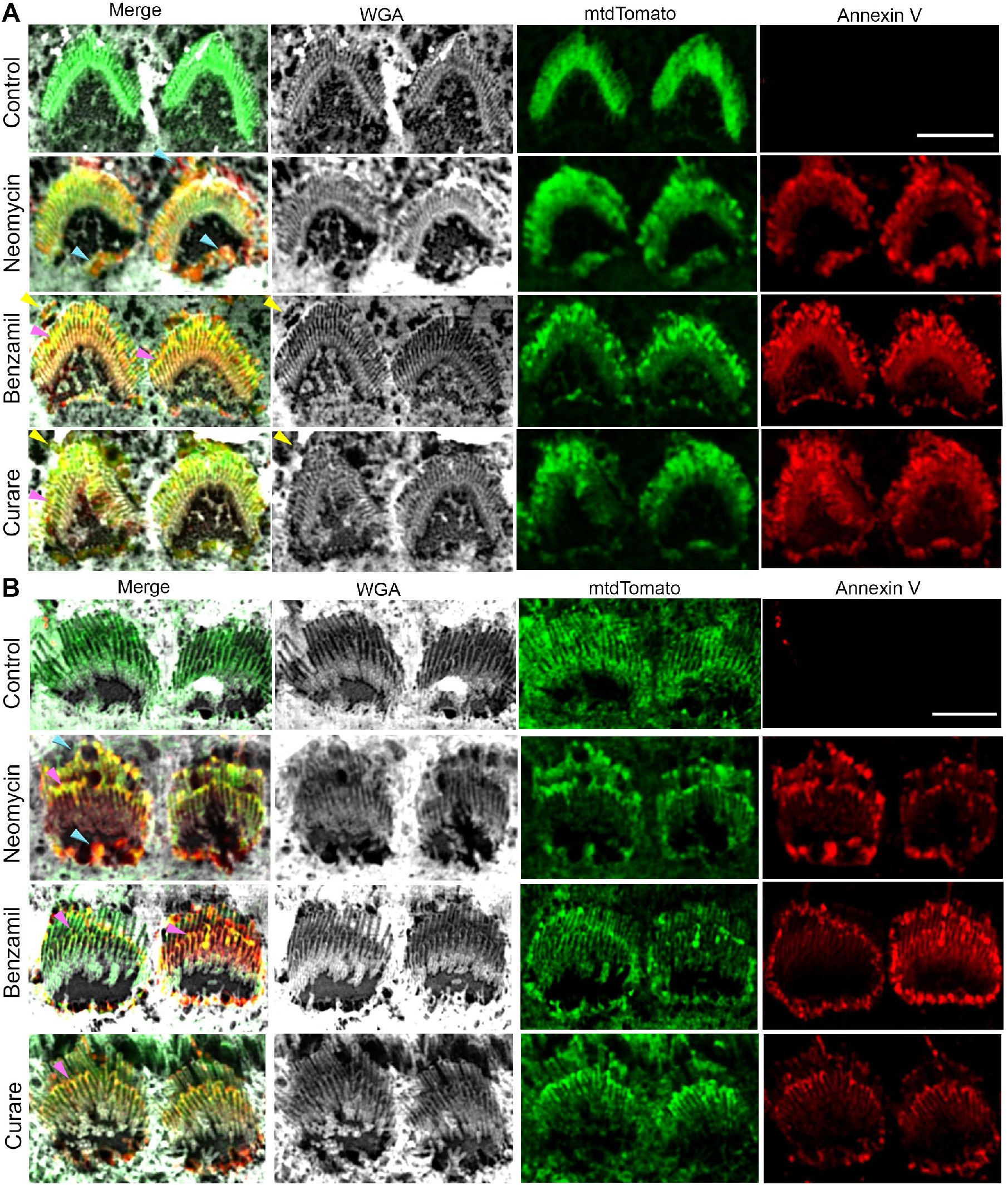
Localization of externalized PS and the mtdTomato reporter in P6 mT/mG^Tg/+^ OHCs and IHCs. Zoom-in confocal images of OHCs (**A**) and IHCs (**B**) from the experiments shown in Figure 1C. Yellow arrows indicate WGA, mtdTomato and annexin V positive blebs detaching from the stereocilia. Blue arrows indicate larger blebs at the kinocilium site and around the apical hair cell region. Pink arrows indicate externalized PS and mtdTomato enrichment at the shorter stereocilia rows. Scale bar is 5 μm. Related to Figure 1.

**Figure S2.**
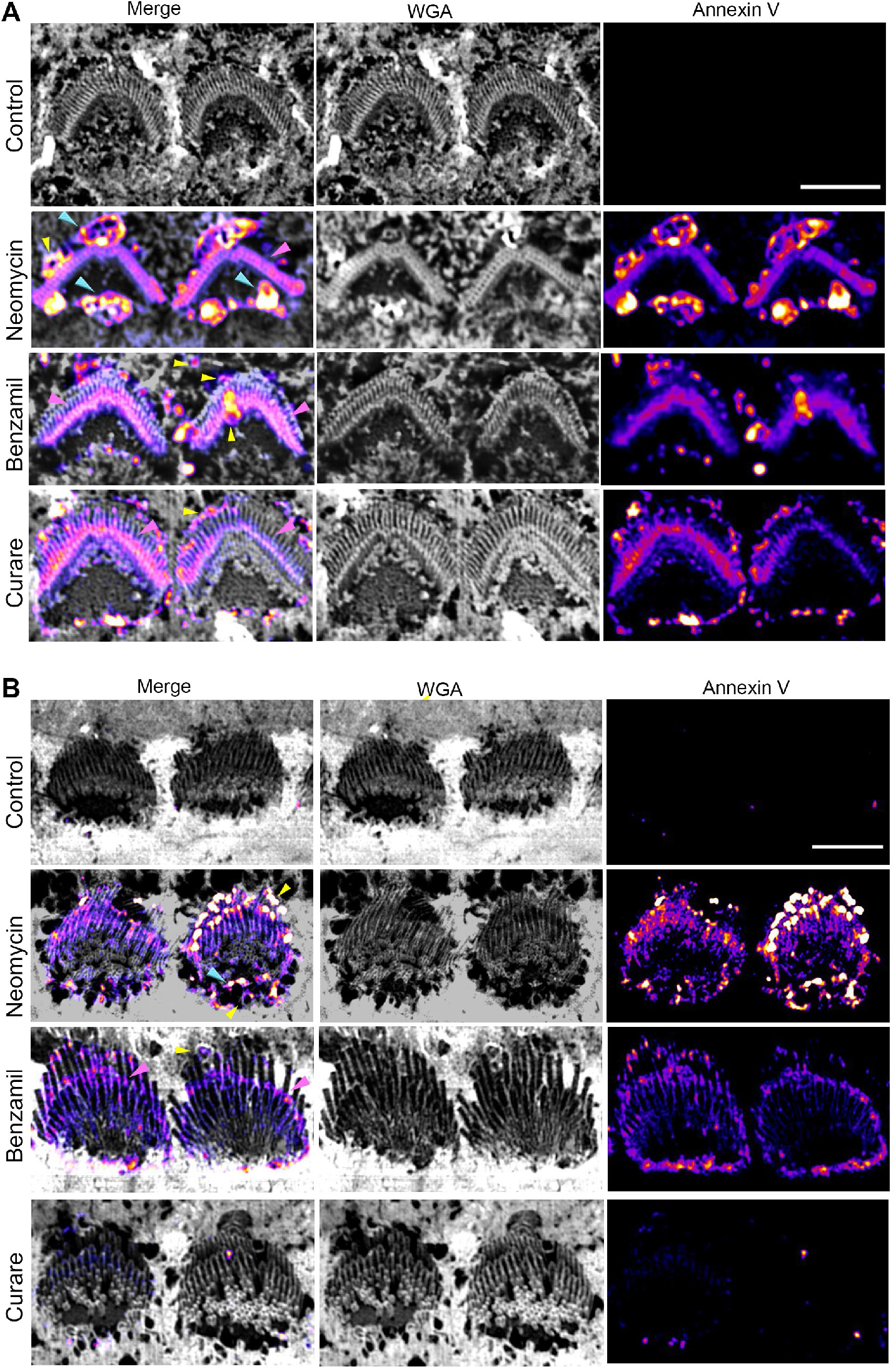
Localization of externalized PS in P6 wild type OHCs and IHCs. Zoom-in confocal images of OHCs (**A**) and IHCs (**B**) from the experiments shown in Figure 1A. Yellow arrows indicate WGA and annexin V positive blebs detaching from the stereocilia. Blue arrows indicate larger blebs at the kinocilium site and around the apical hair cell region. Pink arrows indicate externalized PS at the shorter stereocilia rows. Related to Figure 1.

**Figure S3.**
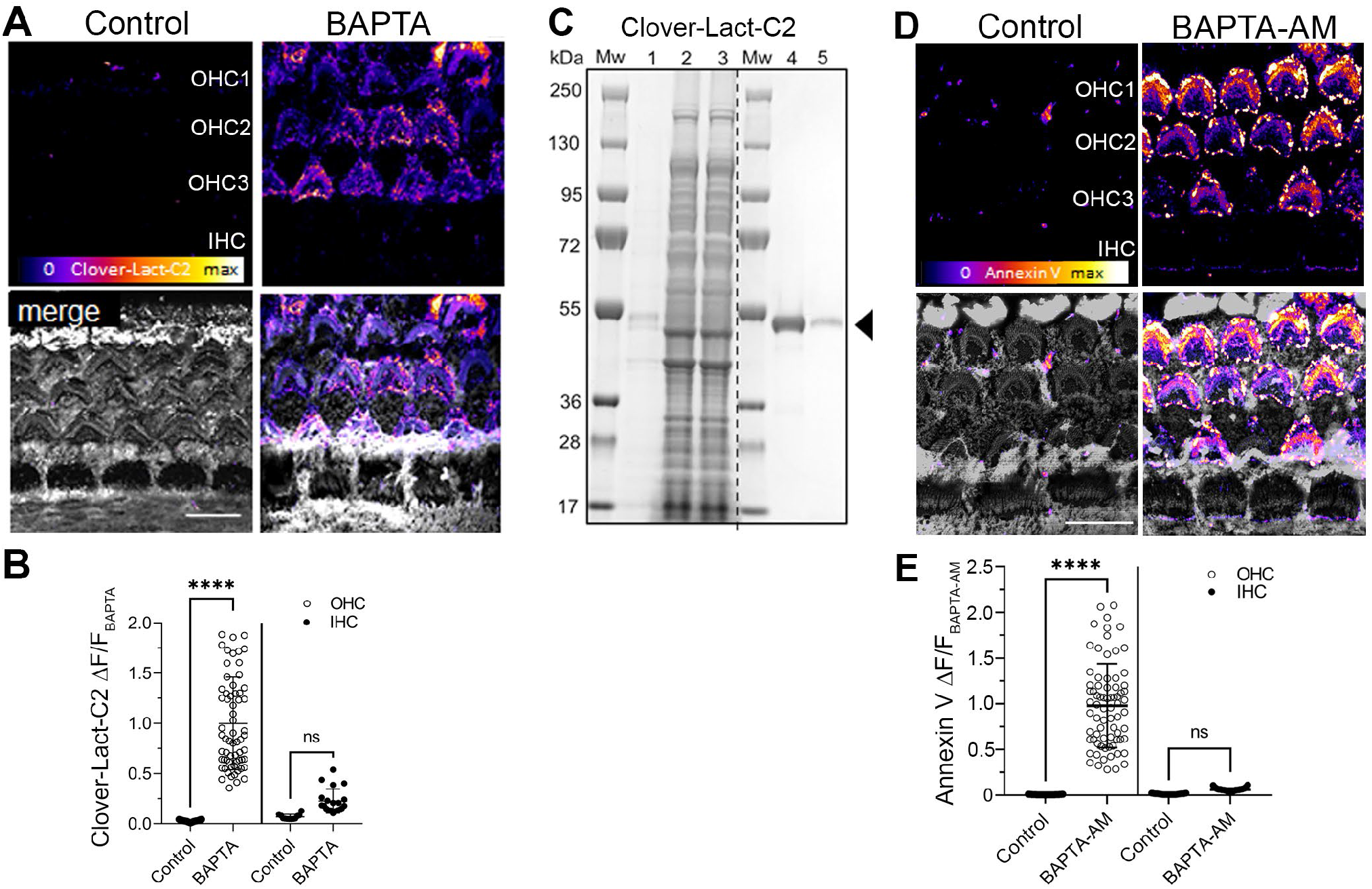
Tip link disruption and buffering of the intracellular Ca^2+^ leads to PS externalization in P6 wild type auditory hair cells. **A)** Related to Figure 2D. Confocal images of hair cells treated with HBSS (control) or 5mM BAPTA in cation-free HBSS buffer (BAPTA) containing Clover-Lact-C2-GFP and WGA (grey). Scale bar is 20 μm. **B**) Quantification of the Clover-Lact-C2-GFP fluorescence intensity at the apical region of IHCs (○) and OHCs (●) treated as in A. **C)** Purification of the 47.2 kDa Clover fluorescent protein followed by a hexa-histidine tag and the C2 domain of lactadherin (clover-Lact-C2). Nupage 4-12% Bis-Tris SDS page protein gel stained with Coomassie showing the flow thoughts (1-3) of the three consecutive washes with 5 mM imidazole in PBS. Line 4 and 5 show two fractions of clover-Lact-C2 protein eluted from the TALON resin with 100 mM imidazole. **D)** Related to Figure 5. P6 wild type hair cells incubated in HBSS containing annexin V in the presence of BAPTA-AM or the vehicle. The scale bar is 10 μm. **E**) Quantification of the annexin V fluorescence intensity at the apical region of the OHCs (○) and IHCs (●) treated as in A. Mean fluorescence intensity ± SD is represented. One-way ANOVA analysis was performed. n.s. p>0.05 and \*\*\*\**p*<0.0001.

**Figure S4.**
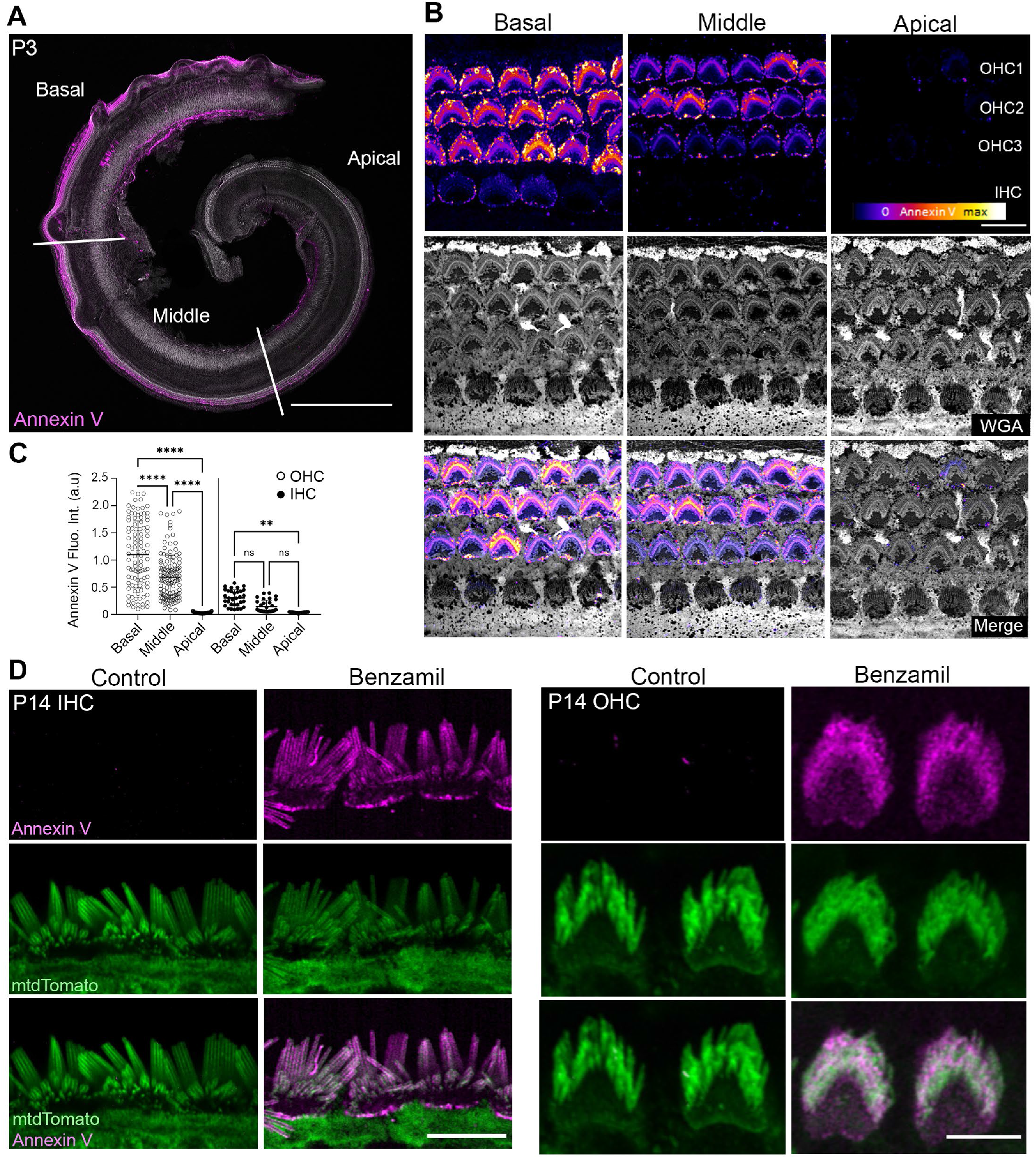
TMC1, but not TMC2, is required for phosphatidylserine externalization and membrane homeostasis. Representative confocal images of TMC1^-/-^ TMC2^-/-^ (**A**), TMC1^-/-^ (**B**) and TMC2^-/-^ (**C**) auditory hair cells incubated with HBSS containing annexin V in the absence (control) or the presence of 100 μM neomycin, benzamil or curare. Annexin V is shown independently (first row) and merged with WGA shown in grey (last row). Outer hair cells (OHC) and inner hair cells (IHC) are indicated. The scale bar is 10 μm. **B**) Quantification of the annexin V fluorescence intensity at the apical region of the OHCs (○) and IHCs (●) in hair cells treated as in A-C. Mean ±SD is shown for n=50-100 hair cells from two cochleae. One-way ANOVA analysis was performed. n.s. p>0.05 and \*\*\*\**p*<0.0001. Related to Figure 3A, 4A and 4D.

**Figure S5.**
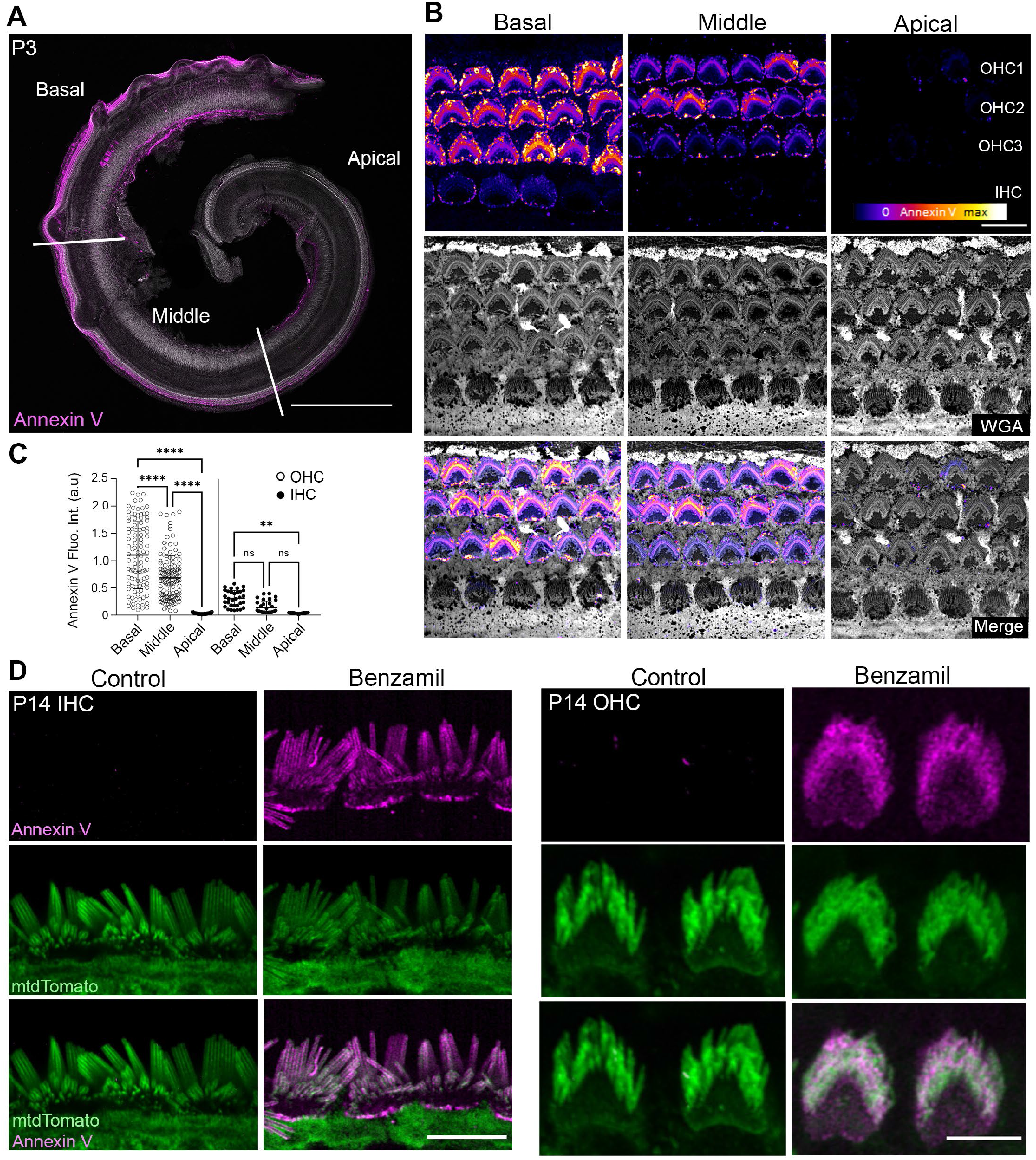
PS externalization follows the tonotopic gradient of the cochlea and is preserved in mature hair cells. **A)** Confocal image of a whole organ of Corti from P3 wild type mice incubated in HBSS containing annexin V and 0.1 mM of curare. The basal, middle and apical regions of the cochlea are indicted. Scale bar corresponds to 500 μm. **B)** Representative confocal images of hair cells at the basal, middle and apical cochlear region are shown for WGA (grey) and annexin V (Fire LUT) channels independently and merged (last row). Outer hair cells (OHC) and inner hair cells (IHC) are indicated. Scale bar is 10 μm. **C)** Quantification of the annexin V fluorescence intensity at the basal, middle and apical region of the OHCs (○) and IHCs (●) in organ of Corti explants treated as in A and B. Mean ± SD is shown. **D**) Confocal image of IHCs and OHCs from P14 mT/mG^Tg/+^ mice expressing the mtdTomato reporter (green) incubated in HBSS containing annexin V (magenta) on the absence (control) or presence of 0.1 mM benzamil. Scale bar is 10 μm.

**Figure S6.**
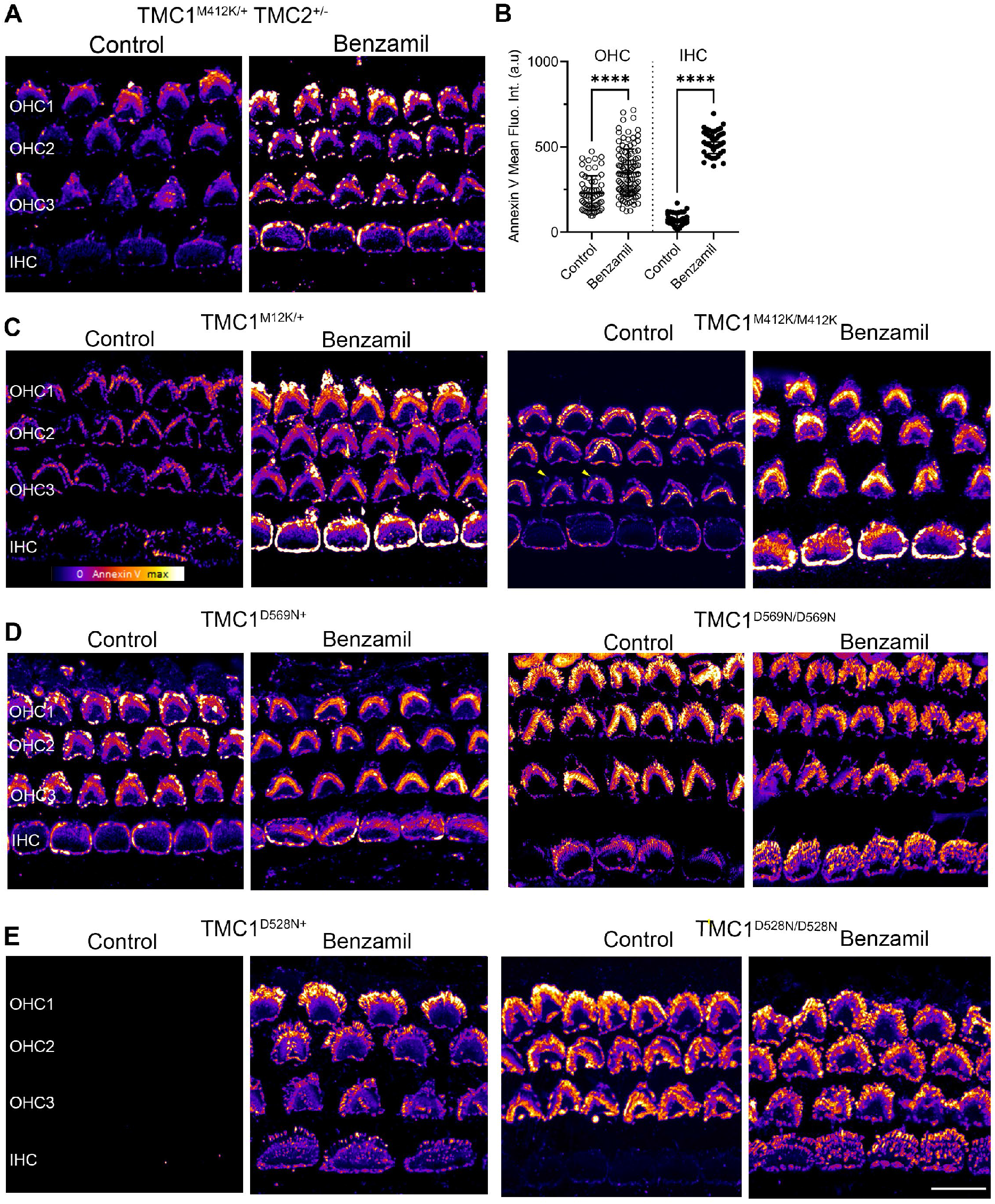
TMC1 deafness mutants present constitutively externalized phosphatidylserine. **A)** Representative confocal images of P6 TMC1^Bth/+^ TMC2^+/-^ hair cells in HBSS containing annexin V in the absence (control) or the presence of 0.1 mM benzamil. **B**) Mean annexin V fluorescence intensity ± SD from one representative cochlea showed in Figure 6. Each dot represents the mean fluorescence intensity of >100 OHCs or >30 OHCs. One-way ANOVA analysis was performed (n.s. p>0.05, \*\**p*<0.01 and \*\*\*\**p*<0.0001). **C-E**) Zoom-out confocal images of TMC1 M412K, D569N and D528N mutant hair cells from the experiments shown in Figure 6. Scale bar is 10 μm. Related to Figure 6.

**FIGURE S7.**
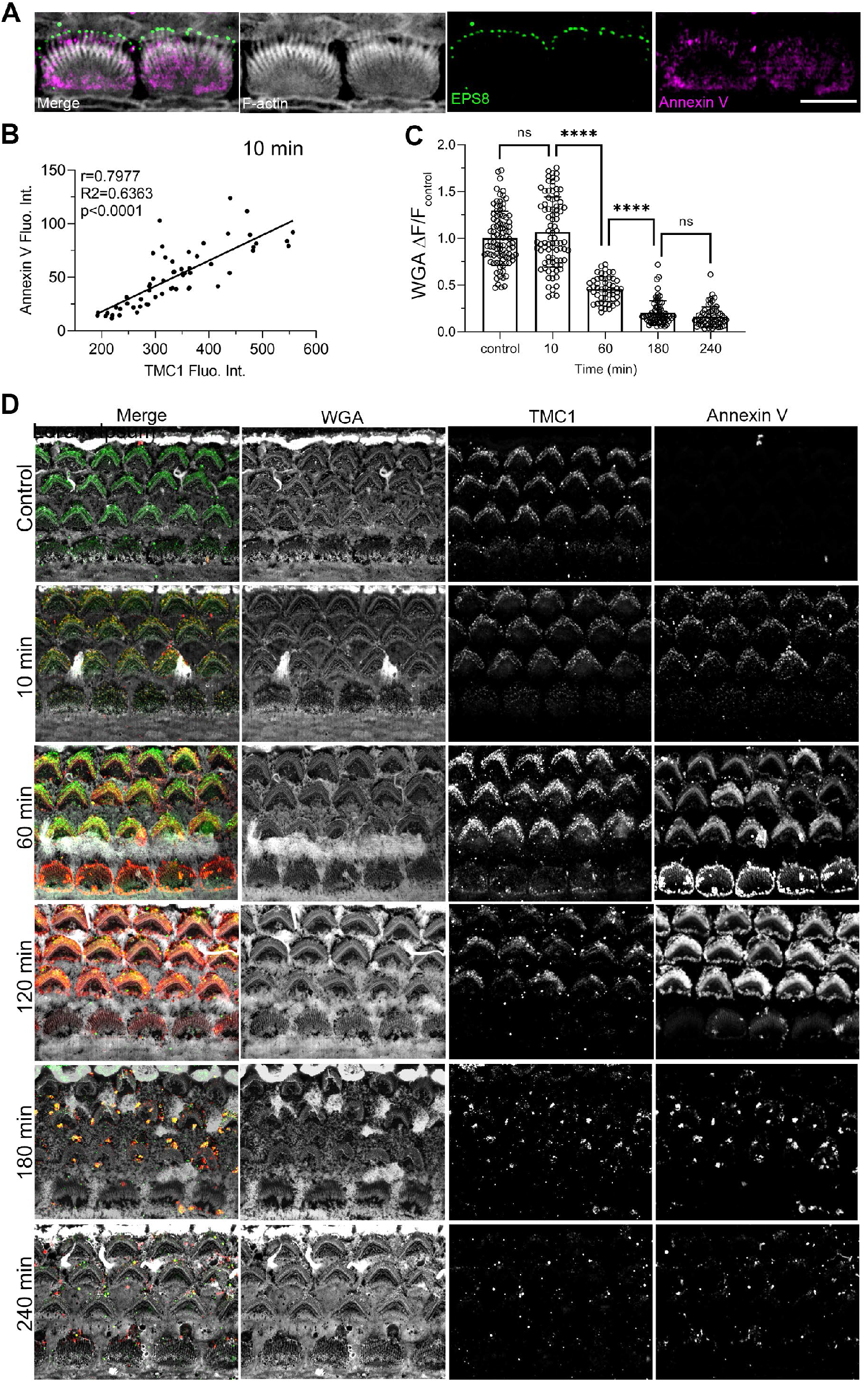
MET blockage and PS externalization alter TMC1 localization at the stereocilia tips. **A)** Annexin V (magenta) positive patches do not localize with ESP8 (green) at the longest stereocilia row of IHCs treated with benzamil. Scale bar is 5 μm. **B)** Correlation between TMC1 and annexin V fluorescence intensity at the apical hair cell region after 10 min incubation with benzamil (n=50-70 OHCs). **C**) Quantification of the WGA fluorescence intensity at the OHCs apical region over time. **D)** Representative zoom-out confocal images of P6 TMC1-cherry mice hair cells non untreated (control) or treated with 100 μM benzamil for 10, 60, 120, 120, and 240 min showed in Figure 7. TMC1-cherry (green) was detected using an anti-RFP antibody, annexin V was used to detect externalized PS (red), and WGA (grey) was included to label the cell surface. Related to Figure 7.

